# Sex and breeding stage differences in neurogenomic profiles reflect hormone signaling in a socially polyandrous shorebird

**DOI:** 10.64898/2026.03.10.710941

**Authors:** Tessa Patton, Evan J. Buck, Aaron Buechlein, Brian W. Davis, Austen J. Ehrie, Erik D. Enbody, Elizabeth M. George, Clemens Kuepper, Jasmine L. Loveland, Leilton W. Luna, Douglas B. Rusch, Quinn K. Thomas, Kimberly A. Rosvall, Sara Lipshutz

## Abstract

In ‘sex-role reversed’ species, females are socially polyandrous and compete for multiple mates, whereas males conduct the majority of parental care. To understand the extent to which physiological differences between females and males are shaped by sex roles, we examined sex differences in gene expression in ‘sex-role reversed’ northern jacanas (Jacana spinosa).

Given that females compete for mating opportunities, and males cycle between courtship and parental care, we predicted that transcriptomic profiles would be more similar between females and courting males, in contrast to female and parenting males. Leveraging a high quality de novo genome assembly, we conducted RNA-seq on two brain regions associated with the regulation of social behavior: the preoptic area of the hypothalamus and the nucleus taeniae. The majority of genes differentially expressed between the sexes were male-biased. Of these male-biased genes, the majority were located on the Z-chromosome. Contrary to our prediction, the greatest difference in autosomal gene expression was between females and courting males, in the preoptic area of the hypothalamus. Several differentially expressed genes related to elements of hormone signaling that are likely to be behaviorally salient, including higher expression of androgen receptor in females relative to parenting males, and higher expression of prolactin receptor in males, regardless of breeding stage. Some sex-associated gene networks were also associated with competitive traits, whereas others were associated with aggressive behaviors, regardless of sex. Few genes were differentially expressed between courting and parenting males, yet some nonetheless had connections to behavioral endocrinology, including prolactin, thyroid and insulin-like growth factor pathways. Our investigation of sex differences in gene expression can help to reveal the molecular mechanisms underlying female competition and male parental care in socially polyandrous species. We conclude that social polyandry is not a simple reversal in the direction of sex-biased gene expression in the brain, but rather a result of complex genetic and hormonal interactions that warrants further study.

## INTRODUCTION

Sex-specific variation in behaviors such as territorial aggression and parental care is widespread across the animal kingdom (Janicke et al. 2016; Zilkha et al. 2017). These behaviors have important fitness consequences, including mating success and offspring survival. When these sexually dimorphic behaviors occur surrounding reproduction, they are often termed sex roles. ‘Conventional’ sex roles are predominantly categorized as male competition over mate acquisition, whereas females are choosy in their mating partners and provide most or all parental care (Bateman 1948; Trivers 1972), but see (Ah-King 2013). In contrast, social polyandry, also known as ‘sex-role reversal,’ is a mating system characterized by female competition for mates and male parental care (Emlen and Oring 1977). Socially polyandrous females compete with each other to mate with multiple males throughout the breeding season and are often larger and more ornamented than males, whereas males conduct the majority of parental care - incubating developing eggs and rearing offspring post-hatching (Emlen and Oring 1977). Given that socially polyandrous systems decouple sex from ‘conventional’ behaviors of competition and parental care, they provide an ideal system to evaluate whether the fundamental mechanisms shaping behavioral profiles reflect sex or sex roles.

Drawing from past research across species with diverse mating systems, there is good evidence that elements of testosterone signaling may be involved in competition for territories or mates in both males (Wingfield et al. 1990) and females (Rosvall et al. 2020). Consequently, modulation of testosterone has been a mechanism of interest for female competition in socially polyandrous species (Lipshutz and Rosvall 2020a). Some studies have found positive correlations between testosterone and competitive traits, for instance the throat patch of the barred buttonquail *Turnix suscitator* (Muck and Goymann 2011) and the wing spurs of female northern jacanas *Jacana spinosa* (Lipshutz and Rosvall 2020b). Other studies found that the gene encoding androgen receptor (*AR*), which binds testosterone in behaviorally salient regions of the brain, had higher expression in females than males in barred buttonquails (Voigt 2016) and black coucals *Centropus grillii* (Voigt and Goymann 2007). These findings suggest that the competitive behaviors observed in social polyandry may stem in part from differential expression of genes encoding receptors of testosterone or other sex steroids in the brain, similar to those found in socially monogamous species (Rosvall et al. 2012; Horton et al. 2014; Munley et al. 2022)

Many studies examining aggression and parental care have focused on the brain, especially the social behavior network, a set of well-conserved regions in the vertebrate brain (Goodson 2005). Several studies of socially polyandrous species have examined brain gene expression in association with female competition and male parental care. For example, the cichlid fish *Julidochromis marlieri* has female-biased dominance and male-biased parental care, whereas its congener *J. transcriptus* has male-biased dominance; critically, a comparative microarray study found shared patterns of brain gene expression in *J. marlieri* females and *J. transcriptus* males, indicating that a conserved set of genes regulates aggressive phenotypes independent of gonadal sex (Schumer et al. 2011). Another study of whole brains in polyandrous pipefish, *Syngnathus scovellli*, found that pregnant males had higher expression of *PAQR6*, a receptor that binds progestin, which is a hormone involved in mammalian pregnancy (Beal et al. 2018). Transcriptome-wide analyses also may shed light on sex-specific selection pressures, including the intensity of sexual selection and sexual conflict (Grath and Parsch 2016; Tosto et al. 2023). With socially polyandrous species, we can therefore gain insights on how sexual selection shapes transcriptomic evolution by investigating the number of genes with sex-biased expression, as well as using co-expression network approaches to connect transcriptomic profiles with phenotypically relevant variation (Hudson et al. 2009). These insights can be further deepened when accounting for chromosomal considerations that may predispose certain genes towards higher expression in males vs females. Because of their ZW sex chromosome system and incomplete dosage compensation, birds generally have higher expression of genes located on the Z chromosome in males than in females (Itoh 2007; Mank 2013).

Much work in socially polyandrous species has focused on sex differences in only one breeding stage, but sex differences may vary temporally across reproductive cycles (Kohl et al. 2017). The transition from territorial establishment to parental care involves substantial shifts in behavior, facilitated by dynamic changes in hormone secretion, gene expression, and neural plasticity (Bentz et al. 2019; Bukhari et al. 2019; Feldman et al. 2019). However, discerning whether differences in gene expression are attributable to sex or sex-associated behaviours requires studying both sexes, not just one. Including both sexes in the same study allows for a more comprehensive understanding (Smiley et al. 2024) and will better isolate how the degree of sex differences shifts across different reproductive stages of competition and parental care.

Here we investigate whether sexual dimorphism in competition and parental care is reflected in the neurogenomic profiles of the northern jacana (*Jacana spinosa*), a species in which females compete for mates and males provide uniparental care for offspring (Jenni and Collier 1972). We generated a state-of-the-art reference genome that we annotated and assembled to the chromosome level. Using this we conducted the first genome-wide study of gene expression in a socially polyandrous bird. Focused on two regions of the social behavior network in the brain, the preoptic area of the hypothalamus and the nucleus taeniae, we compared sex- and stage-specific patterns of 1) differential gene expression and 2) co-expression with other sex-associated phenotypes. Due to the shared competitive state between females and courting males, we hypothesize that females will have more similar neurogenomic profiles to courting males than to parenting males, indicating that sex differences in the transcriptome can help explain sex-specific divergence in behaviors.

## METHODS

### Study population

We conducted field work in La Barqueta, Chiriqui, Panama (8.207N, 82.579W) from 4 June to 9 July 2018. We observed individuals for several hours to confirm their territorial, paired, and reproductive status. We mapped territories relative to geographic landmarks along water canals, small ponds, and irrigated fields. In jacanas, female territories encompass the territories of multiple male mates (Jenni and Collier 1972). Therefore, we avoided sampling adjacent male territories for which we could not distinguish the territory holders. We observed whether females were paired (n = 12), whether males were courting based on copulatory behavior (n = 5), or incubating based on their attending to a nest with eggs (n = 7). These sample sizes are limited, in part due to the logistical challenges of quickly capturing targeted individuals in a tropical, wetland habitat. We later confirmed reproductive status upon collection (details below). Some data from these same individuals has already been included in publications, including testosterone and competitive traits (Lipshutz and Rosvall 2020b), sperm morphology (Lipshutz et al. 2023), and microbiome diversity (Houtz et al. 2025).

### Measuring aggressive behaviors

We measured aggression using a short (5-minute) resident intruder assay, as in Lipshutz & Rosvall (2020b), and we include these data here to relate gene expression to behavior. Briefly, we made moveable taxidermy decoys from females in an aggressive posture and played conspecific calls from a Bluetooth speaker. We rotated through unique combinations from four different decoys and four different vocal stimuli. We placed the decoy and speaker in the center of the focal male and/or female’s territory. We then observed behavior of the territorial female and/or male over the 5-minute trial, narrating into a voice recorder. Aggressive behaviors included average distance from the decoy as well as number of flyovers, swoops, wingspreads, and vocalizations. All simulated territorial intrusions were conducted between 9am and 6pm.

### Measuring morphological traits

We measured several morphological traits hypothesized to facilitate mating competition in jacanas: wing spur length (mm), body mass (g), and tarsus length (mm). Previous analyses of these same individuals found positive correlations between testosterone and wing spur length in females, as well as testosterone and testes mass in males (Lipshutz and Rosvall 2020b); we include these morphological data again here to relate these competitive traits to gene expression. Notably, wing spurs are sharp keratinous sheaths over metacarpal bone growth that jacanas use as weapons and signals during aggressive posturing. Wing spurs, body mass, facial shields, and tarsi are larger in females than in males (Lipshutz 2017). In a congener, the wattled jacana (*Jacana jacana*), both female and male territory holders had larger wing spurs, body mass, facial shields, and tarsi than non-territorial floaters, indicating the role of these traits in mediating competition for territories and reproductive success (Emlen and Wrege 2004).

### Sample collection

Our goal was to quantify gene expression in the brain in a constitutive, naturalistic state.

All birds were collected using an air rifle, followed by anesthetic overdose of isoflurane and decapitation. Some individuals (n = 12) were collected quickly after the short aggression assay (average time from assay start to euthanasia = 9 min 20s ±1 min 35s). Because transcription of immediate early genes after introduction of a social stimulus takes at least 30 min to show a transient increase in expression (Burmeister et al. 2005), we assume that measures of gene expression represent constitutive levels. For individuals that could not be collected immediately, we returned to territories 5-8 days later for collection (n = 6 females, 6 males). For these ‘delayed’ collections, we monitored with daily behavioral observations to ensure that male breeding stage did not change. We found no difference in testosterone levels between immediate vs. delayed samples for either females or males (Lipshutz and Rosvall 2020b), so we combined these immediate vs. delayed collection individuals for further analysis.

We collected trunk blood for testosterone quantification, published in (Lipshutz and Rosvall 2020b). We include these testosterone data here to examine how they relate to gene expression. Trunk blood was centrifuged, plasma was drawn off, and samples were frozen in liquid nitrogen followed by a −20°C freezer until later quantification using enzyme immunoassay. Using RNAse-free tools, we dissected whole brains from the skull, flash froze in powdered dry ice, and stored in dry ice and then a −80°C freezer until further processing (average time from euthanasia to brain tissue collection = 7 min 10s ± 20s). All individuals were in breeding condition, as evidenced by yolky, developed follicles for females, enlarged testes for courting males, brood patches for incubating males, and testosterone profiles that differed by sex and breeding stage (Lipshutz and Rosvall 2020b).

### Brain microdissection and RNA isolation

To isolate RNA from focal brain regions, we used a CM1850 cryostat (Leica Biosystems, Buffalo Grove, IL, USA), and sectioned along the coronal plane (as defined by (Puelles 2007)), preceding from caudal to rostral. Because this is the first brain study for any species in the family Jacanidae, we first created a series of 40uM brain sections stained with cresyl violet and scanned using a Motic EasyScan (Motic, Richmond, British Colombia, CA) to aid in identifying anatomical landmarks and focal regions from brain atlases (see below). For the remaining RNA samples, we sectioned brain tissue at 200 uM. Cryostat temperature ranged from -14 to -16°C.

We thaw-mounted sections for <10 sec onto glass slides (Fisher superfrost 12-544-2, uncharged) and then immediately refroze them at -43°C using the quick-freeze shelf within the cryostat chamber. We stored slides for up to a week at -80°C before microdissection.

Our microdissection methods followed (Horton et al. 2020). We set up a Styrofoam cooler with a bed of dry ice and placed a chilled marble slab and coring mat on top. We used an LED magnifying lamp to locate brain regions using anatomical landmarks (e.g. ventricles, fiber tracts). We collected neural tissue with rapid-core reusable biopsy punches (World Precision Instruments, Sarasota, FL, USA). Our punch tool size varied based on focal region (details below). We expelled tissue cores into microcentrifuge tubes on dry ice and kept them frozen until all samples were dissected for that subject. We then added TRIzol (Invitrogen, Carlsbad, CA, USA) directly to the tubes, briefly vortexed, and stored at -80°C until RNA extraction (up to 5 months).

We focus on two conserved regions of the brain associated with agonistic behavior and parental care in other taxa: the nucleus taeniae, the avian homologue of the medial amygdala, and the preoptic area of the hypothalamus (Goodson 2005). For the nucleus taeniae, we used neuroanatomical landmarks following the chick atlas (AHi subnuclei and ATn, Puelles, 2007) and the ruff atlas (Loveland et al. 2025). Our preoptic area of the hypothalamus samples contain partial sampling of the anterior and medial preoptic area (AMPO and MPO, according to Puelles, 2007), anterior hypothalamus (see plate 8.2 in Kuenzel and Masson 1988), and periventricular hypothalamic nucleus (PHN, (Kuenzel and Golden 2006)). For the nucleus taeniae, we used a 0.75 mm bilateral punch on both hemispheres, followed by 0.5 mm for the last two sections (Figure S1). For the preoptic area of the hypothalamus, we used a 1.2 mm single punch (in the middle). We gave each tissue core a confidence score from 1-5 based on the accuracy of the punch location, and all cores included in this study scored 3 or higher.

We extracted total RNA from tissues following the phenol-chloroform-based TRIzol manufacturer protocol, adjusted for nano-scale brain tissue volumes, and with the addition of phase-lock gel tubes (QuantaBio) and GlycoBlue (Invitrogen, Carlsbad, CA, USA) to facilitate pellet precipitation. We resuspended RNA in nuclease-free, Ultrapure water (Invitrogen). We analyzed RNA quantity and quality using a TapeStation 4150 and high-sensitivity RNA screentape (Agilent Technologies). Mean sample RIN for the preoptic area of the hypothalamus was 8.9 ± 0.2 and for nucleus taeniae was 8.7 ± 0.1. Mean sample concentration for the preoptic area of the hypothalamus was 46.8 ± 4.5 ng/µl and for nucleus taeniae was 59.7 ± 7.5 ng/µl.

### RNA-seq and mapping

We submitted total extracted RNA to Indiana University’s Center for Genomics and Bioinformatics for sequencing. We constructed complementary DNA libraries using a TruSeq Stranded mRNA HT Sample Prep Kit following the manufacturer’s protocol. Sequencing was performed using an Illumina NextSeq 500 platform with a 75-cycle sequencing module generating 38-bp paired-end reads. We generated an average of ∼28 million reads for the barcoded RNA samples (Table S1).

We demultiplexed sequencing reads with bcl2fastq (v2.20.0.422). We cleaned reads with Trimmomatic v0.39 (Bolger et al. 2014) and mapped reads to the jacana genome (see below) using Hisat2 v2.2.1 (Kim et al. 2019). We only included reads mapped in proper pairs, sorted and indexed using Samtools v1.19.2 (Li et al. 2009). For nucleus taeniae samples, ∼21 million read pairs per sample were mapped, which account for ∼82% of the total trimmed read pairs (Table S1). For preoptic area of the hypothalamus samples, ∼22 million read pairs were mapped, which account for ∼85% of the total trimmed read pairs (Table S1). Reads were quantified into genes using featureCounts from the SUBREAD package v2.0.0 (Liao et al. 2019).

### HiFi reference genome and HiC chromosome-level assembly

Genomic resources were generated from two females, of the same individuals included in this study. For one female (NOF2), we generated a reference genome using the PacBio HiFi (Pacific Biosciences, Menlo Park, CA) services at Maryland Genomics. They conducted a high molecular weight gDNA extraction from approximately 25 milligrams of ethanol-preserved Jacana spleen tissue. The tissue underwent an ethanol removal procedure prior to processing with the standard Dounce homogenizer protocol for tissue via the Circulomics Nanobind DNA Kit. The tissue was homogenized and lysed. The lysate was then bound to a magnetic Nanobind disk to harvest genomic DNA. The resulting genomic DNA was assessed via the Femto Pulse instrument (Agilent, Santa Clara, CA) to confirm its integrity prior to library preparation.

Ten micrograms of genomic DNA sample were sheared on MegaRuptor 2 (Diagenode, Denville, NJ), at 15 kb setting. Five micrograms of sheared genomic DNA were prepared into a PacBio sequencing library using SMRTbell Express Template Prep Kit 2.0 (Pacific Biosciences, Menlo Park, CA) following manufacturer’s instructions. After library preparation, the library was size-selected on the BluePippin instrument (Sage Science, Beverly, MA) to remove library fragments smaller than 10 kb. Prior to sequencing, the library was bound to sequencing polymerase with the Sequel II Binding kit 2.2, then sequenced with the Sequel II Sequencing kit 2.0 and SMRT cell 8M on the Sequel II instrument (Pacific Biosciences).

For a second female (NOF7), flash-frozen spleen tissue was prepared for Hi-C at Texas A&M University to identify chromatin interactions and scaffold chromosomes. We assembled the northern jacana genome *de novo* using Hifiasm v0.16.0, a haplotype-resolved assembler for PacBio HiFi reads (Cheng et al. 2021), in combination with Hi-C data, generating 663 scaffolds. These scaffolds were then ordered and oriented using minimap2 v.2.30 (Li 2018) and RagTag v2.1.0 (Alonge et al. 2022), with chicken (*Gallus gallus*) and zebra finch (*Taeniopygia guttata*) genomes as references. The chicken genome provided better synteny results and was therefore used as the main guide. The final chromosome-level assembly of the jacana genome consists of 41 mapped chromosomes, including both Z and W, with approximately 95% of the scaffolds successfully placed onto chromosomes. This chromosome number aligns with a karyotype from a congener, the wattled jacana (*J. jacana*) (Kretschmer et al. 2020).

To assess gene content completeness, we used BUSCO v4.1.4 (Simão et al., 2015) with the avian ‘aves_odb10’ lineage (Table S2, Figure S2). Gene annotation was performed by lifting over annotations from the chicken (*Gallus gallus)* chromosome-level genome assembly (GRC7b, GCA_016699485.1) available on ensembl.org. To accomplish this, we used Liftoff v1.6.3 (Schumate and Salzberg 2021) to align all chicken genes to the northern jacana scaffolds. Liftoff identifies high-confidence alignments for all exons using minimap2 v.2.30, while taking into account transcript and gene structure. This method does not detect novel genes or novel splice events or exons, but is suitable for RNA-seq mapping. We successfully assigned 14,588 out of 17,007 genes from the chicken genome. The assembled jacana genome is available on NCBI under version JBQSLJ010000000.

### Identifying Z- and W-linked genes

We identified putative Z and W chromosome-linked scaffolds using depth of coverage from whole-genome sequences of 5 female and 5 male northern jacanas, which were different individuals from this gene expression study (Luna and Lipshutz unpublished). First, we aligned and mapped raw reads to scaffolds of the northern jacana genome using BowTie2 (Langmead and Salzberg 2012) with the “very-sensitive-local” preset, setting the maximum fragment length for valid paired-end alignments (-X) to 700 bp. PCR duplicates were then marked using PicardTools v2.20.8 (https://broadinstitute.github.io/picard/). Next, we calculated the scaffold depth of coverage with the SACT package in R (Nursyifa et al. 2022). Normalized coverage depths were used to differentiate heterogametic ZW scaffolds from homogametic autosomal and ZZ scaffolds. To validate the Z-linked and W-linked scaffolds identified through coverage analysis, we mapped these scaffolds to the chicken and zebra finch genomes, following similar mapping guidelines as above. In the case of discrepancies for some gene locations across species, we further confirmed putative W chromosome-linked genes based on their lack of gene expression in male jacanas in our RNAseq dataset. For instance, of genes with female-biased expression in jacanas, MIER3, ZFAND5, RPL37, PIAS2, and VCP are on the Z chromosome in the chicken genome, but their lack of Z-assignment in male jacanas both from whole-genome depth of coverage data, together with lack of transcript expression, suggests they are restricted to the W chromosome in the northern jacana. In total, we identified 534 Z-linked genes and 16 W-linked genes.

### Differential expression analyses

We used DESeq2 v 1.38.3 (Love et al. 2014) to normalize read counts, which we visualized with a PCA (Figure S3). For each brain region, we compared log2fold changes of differential gene expression between three dyads: 1) females and courting males, 2) females and parenting males, and 3) courting and parenting males. For each transcript, we fit count data to a negative binomial generalized linear model with a fixed effect for sex and breeding stage.

We identified significantly differentially expressed transcripts using a per-transcript Wald test statistic. We corrected p-values for multiple-testing using the Benjamini–Hochberg method (adjusted p ≤ .05).

### Weighted Gene Co-expression Network Analyses

To account for the collective, rather than individual, action of genes, we conducted weighted gene co-expression network analyses (WGCNA) for each brain region (Langfelder and Horvath 2008). Sample clustering identified one outlier in the nucleus taeniae analyses, a female, and one outlier in the preoptic area of the hypothalamus analysis, a courting male, which we then excluded from WGCNA. We normalized and filtered out genes with <10 counts in 90% of samples and used the 75% most variable genes (8863 in the nucleus taeniae, 9080 in the preoptic area of the hypothalamus). We generated a signed hybrid network using a soft threshold power (β) = 12 for the nucleus taeniae and (β) = 14 for the preoptic area of the hypothalamus. We used a biweight midcorrelation function and constructed modules with a minimum size of 30. We then merged modules with Dynamic Tree Cut using a threshold size of 0.25, as these genes are highly co-expressed.

To explore potential relationships between brain gene expression and competitive traits, we correlated expression levels for each module’s first eigengene with traits of interest. These traits included male breeding stage, sex, log testosterone, and gonad mass, as well as behavioral responses to the resident-intruder assay, including distance to decoy, wherein shorter distance indicates higher aggression, as well as swoops, vocalizations, and wingspreads. Given that body mass, tarsus length, and wing spur length may be correlated, we calculated a composite size morphology trait using a principal component analysis. Because these morphological traits differ greatly between the sexes and are much larger in females (Lipshutz 2017), we thus interpret modules that correlate strongly with sex *and* size morphology to be driven by sex-confounded differences in both phenotypic traits and gene expression. To evaluate additional confounding variables, we also included Julian date, time from resident intruder assay to collection, and time from collection to brain dissection; none of these metrics were correlated with any trait-associated modules, so we did not consider them further. For each module significantly associated with traits of interest, we examined genes with high module membership (MM>|0.80|) and gene significance (GS>|0.60|), hereafter ‘hub genes’ (Table S8, S9). We also plotted module membership versus gene significance for modules significantly associated with traits of interest (Figure S7-13). In cases where modules were associated with sex, we examined whether there was overrepresentation of genes on the Z chromosome.

### Gene ontology

We inferred gene function based on genecards.org (Stelzer et al. 2016) as well as gene ontology (GO) analyses. For each set of differentially expressed genes, as well as genes from WGCNA modules associated with traits of interest (module membership |>| 0.6) we inferred GO terms of biological processes using PantherGO (Mi et al. 2019). We evaluated gene enrichment using *Homo sapiens* as the reference because its ontologies are orthologous to, but more complete than avian references. We limited our reference list to the 14,588 genes previously identified from our de-novo genome annotation. We considered GO terms with an FDR < 0.05 as significantly enriched. We report the most specific subclass (first descendent) of each significant GO term.

### Tissue specificity

Recognizing the potential for autosomal and Z-linked genes to vary in their expression, and because this is the first transcriptomic study in the family Jacanidae, we conducted a broader comparison across three different tissues to characterize the specificity of gene expression across tissues and among Z-linked genes. From one representative individual per group (female, courting male, parenting male), we extracted RNA and sequenced whole blood and gonad samples. To assess the specificity of expression in the preoptic area of the hypothalamus and nucleus taeniae, we combined the remaining brain tissues from the same individuals, reflecting whole brain gene expression. We assessed the percentage of genes expressed in each tissue transcriptome for all genes, as well as just Z-linked genes (Figures S15, S17). In addition to describing whether genes were expressed in each tissue, we also calculated the tau index of gene tissue sensitivity, to assess how unevenly each gene is expressed across tissues (Figure S16).

## RESULTS

### Sex-biased gene expression

#### Sex-biased gene expression varies by brain region and chromosome

When viewing both brain regions in a PCA, most of the variance among samples was explained by tissue type, then sex and breeding stage (Figure S3). In each brain region, we identified hundreds of differentially expressed genes between females and males of both breeding stages, comprising 3-4% of genes in the transcriptome (Figure 1, Tables S3-7). In the preoptic area of the hypothalamus, autosomal DEGs made up a greater proportion of sex-biased genes between females and courting males than between females and parenting males (Fisher’s exact test, p < 0.001) (Figure 1, Table S3). In contrast, in the nucleus taeniae, the proportion of autosomal DEGs was similar between sex-stage comparisons (Fisher’s exact test, p > 0.05).

The majority of genes differentially expressed between sexes had higher expression in males (83-85%) (Figure 2). Of these male-biased genes, the majority (60-73%) were assigned to the Z chromosome. However, 40-50% of total genes with sex-biased expression were found on autosomes. Of the female-biased genes, most (91-93%) were on autosomes; few (7-9%) were assigned to sex chromosomes (Figure 2). This included 8 genes on the W chromosome (*CHD1W*, *MIER3, PIAS2, RPL37, VCP, ZFAND5*, and two genes without annotation, ENSGALG00010011555 and ENSGALG00010012357) as well as one gene on the Z chromosome (*AQP3*) for females vs parenting males in the preoptic area of the hypothalamus. None of these sex-biased gene lists, whether autosomal, Z-linked, W-linked, or total sex-biased genes, were significantly enriched for any GO terms.

**Figure 1:**
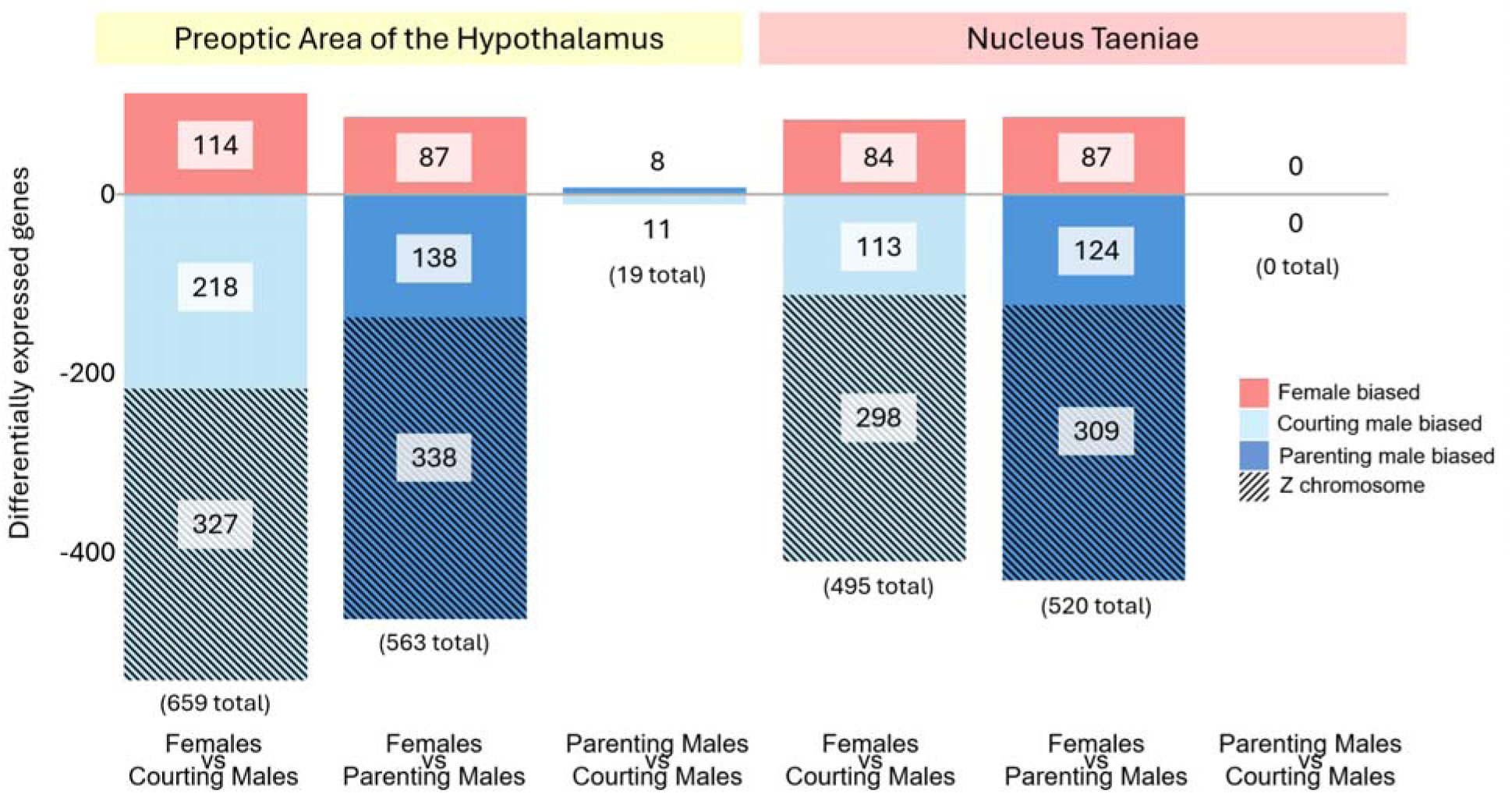
The number of significantly differentially expressed Z-linked and autosomal genes (adjusted p<0.05) for each sex (female, male) and male breeding stage (parenting, courting) comparison in the preoptic area of the hypothalamus and nucleus taeniae. Genes more highly expressed in females (all courting) are shown in red (i.e female biased), genes more highly expressed in courting males are shown in light blue (i.e. courting male biased), and genes more highly expressed in parenting males are shown in dark blue (i.e. parenting male biased). Genes on the Z chromosome are striped. Genes on the W chromosome are shown in Figure 2.

#### Many sex-biased genes relate to hormone and neuropeptide signaling

Despite the lack of GO enrichment among sex-biased genes, several candidate genes are likely to be biologically relevant for sex differences in behavior in this system. On the autosomes, there were several genes associated with hormonal and neuropeptide signaling, and their degree of dimorphism in gene expression varied by brain region and male stage.

Several steroid-related genes had higher expression in females. In the preoptic area of the hypothalamus, androgen receptor (*AR*) had higher expression in females than parenting males, and estrogen receptor 1 (*ESR1)* had higher expression in females than courting males (Figure 3). Estrogen receptor 2 (*ESR2*) and aromatase (*CYP19A1*) were not differentially expressed between sexes in either brain region. In the nucleus taeniae, neuropeptide Y (*NPY*) had higher expression in females than in parenting males. Several other hormone-related genes had highe expression in males. In the preoptic area of the hypothalamus, iodotyrosine deiodinase (*IYD*), which encodes a thyroidal enzyme, and insulin like growth factor binding protein 3 (*IGFBP3*), which is associated with the insulin-like growth factor (*IGF*) pathway, had higher expression in parenting males than females (Figure S4).

Male-biased genes on the Z chromosome tended to be higher in males of both stages in both brain regions (Figure 2). Male-biased, Z-linked genes related to hormone signaling included prostaglandin reductase 1 (*PTGR1*) as well as prostaglandin E receptor 4 (*PTGER4*). The Z-linked prolactin receptor (*PRLR*) had higher expression in males than females (Figure 3), though this sex difference was greater between parenting males than courting males. Additional male-biased, Z-linked genes included StAR-related lipid transfer protein 4 (*STARD4*), a cholesterol transporter, and 5’-AMP-activated protein kinase catalytic subunit alpha-1 (*PRKAA1*), a gene product involved in the gonadotropin-releasing hormone pathway, glial cell line-derived neurotrophic factor (*GDNF*), neurotrophic receptor kinase 2 (*NTRK2*), and rapamycin-insensitive companion of mTOR (*RICTOR*) (Stelzer et al. 2016).

**Figure 2:**
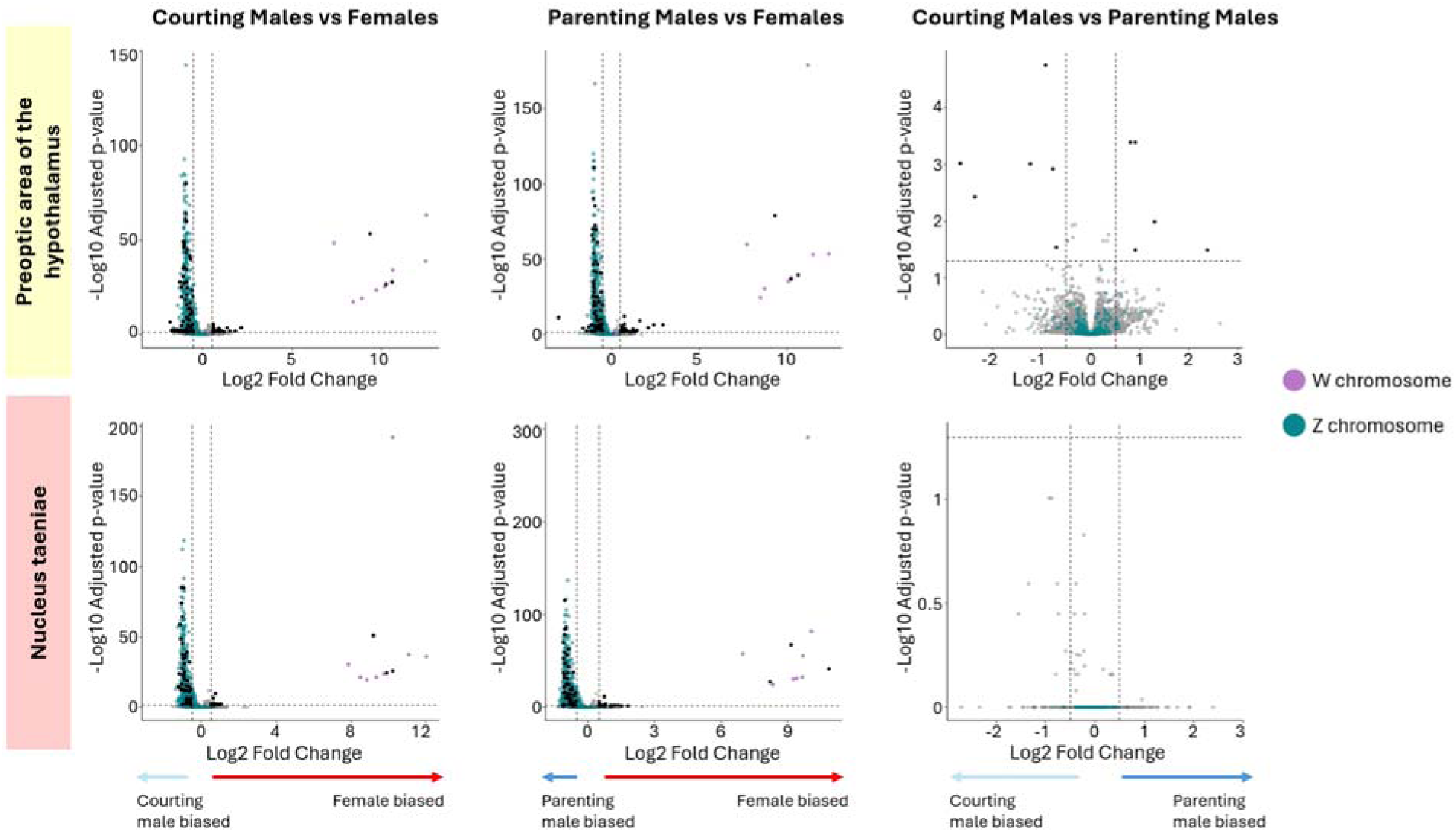
Volcano plots for each sex-stage comparison in each brain region. The -log10 P-value is plotted on the y-axis, with log2-fold change across the x axis. Each dot represents a gene. Colors represent whether the gene was on the Z chromosome (teal), W chromosome (purple), or autosomal (black and grey). Black dots represent autosomal genes that pass the P-value threshold (adjusted p<0.05), the log2-fold change threshold (log2-fold change<|0.5|), or both. Gray dots represent autosomal genes that do not pass the P-value threshold (adjusted p<0.05), the log2-fold change threshold (log2-fold change<|0.5|), or both. The log2-fold change threshold is used for visualization purposes, but not for the significance cutoff. Arrows indicate whether a gene is more highly expressed in females, parenting males, or courting males.

#### *Few* differentially *expressed genes between male breeding stages*

Few genes were differentially expressed between courting and parenting males – only 19 genes in the preoptic area of the hypothalamus and no genes in the nucleus taeniae (Figure 1; Table S5). Some of the top differentially expressed genes with higher expression in courting males included prolactin-releasing hormone (*PRLH*), as well as prodynorphin (*PDYN*). Some additional genes with higher expression in parenting males were associated with the thyroid and insulin-like growth factor (*IGF*) pathways, including insulin-like growth factor binding protein 3 (*IGFBP3*) and pappalysin 2 (*PAPPA2*), and iodotyrosine deiodinase (*IYD*) (Figure S4) (Stelzer et al., 2016). Aromatase (*CYP19A1*) was higher in courting males relative to parenting males (Figure 3), but this difference was not significant after correction for false discovery rate.

**Figure 3.**
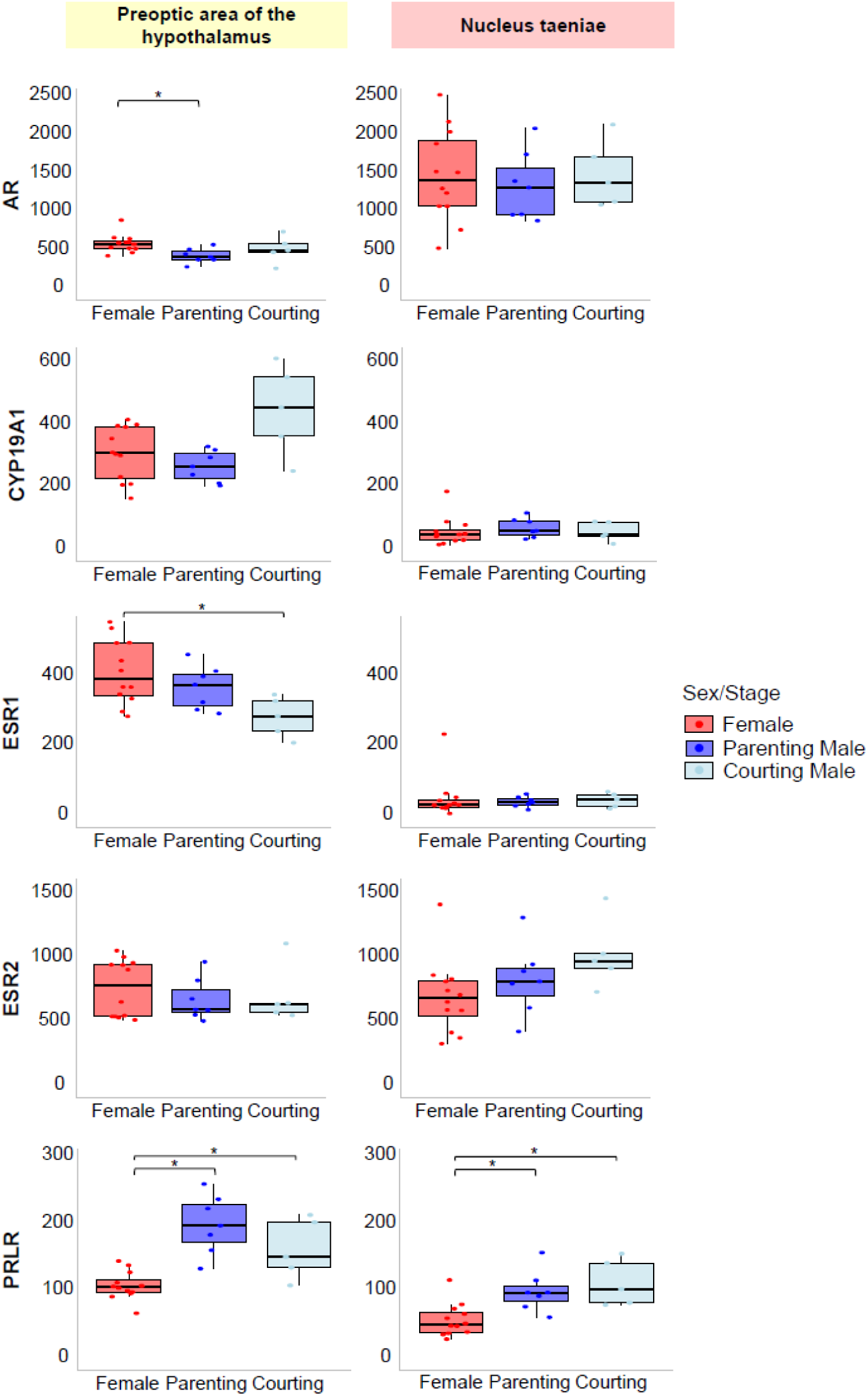
Boxplots of raw count expression for candidate genes associated with hormonal signaling. Boxplots on the left show expression in the preoptic area of the hypothalamus (POA), and boxplots on the right show expression in the nucleus taeniae (TnA). All genes are located on autosomes except PRLR, which is located on the Z chromosome. * indicates p<0.05 after false discovery rate.

### Weighted Gene Co-expression Network Analysis

We identified eight modules of coexpressed genes for the preoptic area of the hypothalamus, and eleven modules for the nucleus taeniae (Figure 3, Table S8, S9). In both brain regions, we identified modules that correlated with sex, male breeding stage, aggression, and morphology PC1. Two modules associated with sex had higher eigengene values in males: the brown module of the preoptic area of the hypothalamus and the blue module of the nucleus taenia. Neither of these sex-associated modules had genes significantly enriched for any GO terms (Table S10). These modules overlapped significantly in gene composition (Fisher’s exact test, p < 0.0001), and nearly 50% of the genes in each of these modules were Z-linked (Figure S14). These modules also negatively correlated with morphology PC1 (body mass, wing spur length, tarsus length), consistent with the fact that these traits were smaller in males. The brown module in the preoptic area of the hypothalamus correlated marginally with aggression (wingspreads), presumably driven by males displaying wingspreads more often than females. In the nucleus taeniae, two modules - greenyellow and cyan - associated with sex had higher eigengene values in females and were positively correlated with morphology PC1 (larger size morphology). This cyan module also included 100% of the nucleus taeniae’s female-biased DEGs (8 putatively W-linked, 3 autosomal), though the module contained 72 genes overall that were not significantly enriched for any GO terms (Table S10). The greenyellow module was enriched for axonogenesis (Table S10).

We also identified several gene modules that were associated with competitive traits, but not with sex (Figure 4), allowing us to better disentangle sex from from some sex-associated traits. In the preoptic area of the hypothalamus, the cyan module was positively correlated with aggression (closer distance to decoy) and was enriched for GO terms including central nervous system myelination and cell-cell adhesion (Table S10). In the nucleus taeniae, the pink module positively correlated with aggression (closer distance to decoy, more vocalizations), and was enriched for GO terms including endocardial cushion to mesenchymal transition, positive regulation of lung ciliated cell differentiation, regulation of cilium beat frequency, and sperm axoneme assembly (Table S10). The nucleus taeniae’s midnightblue module was also positively correlated with aggression (swoops) and significantly enriched for GO terms including protein folding in endoplasmic reticulum, protein refolding, and response to heat (Table S10). Also in the nucleus taeniae, the yellow module correlated negatively with morphology PC1 and was enriched for GO terms including septin cytoskeleton organization and central nervous system myelination (Table S10). Though the yellow module was not significantly correlated with sex, upon closer inspection, its higher expression in females and positive correlation with larger morphology appears confounded by sex differences in size.

**Figure 4:**
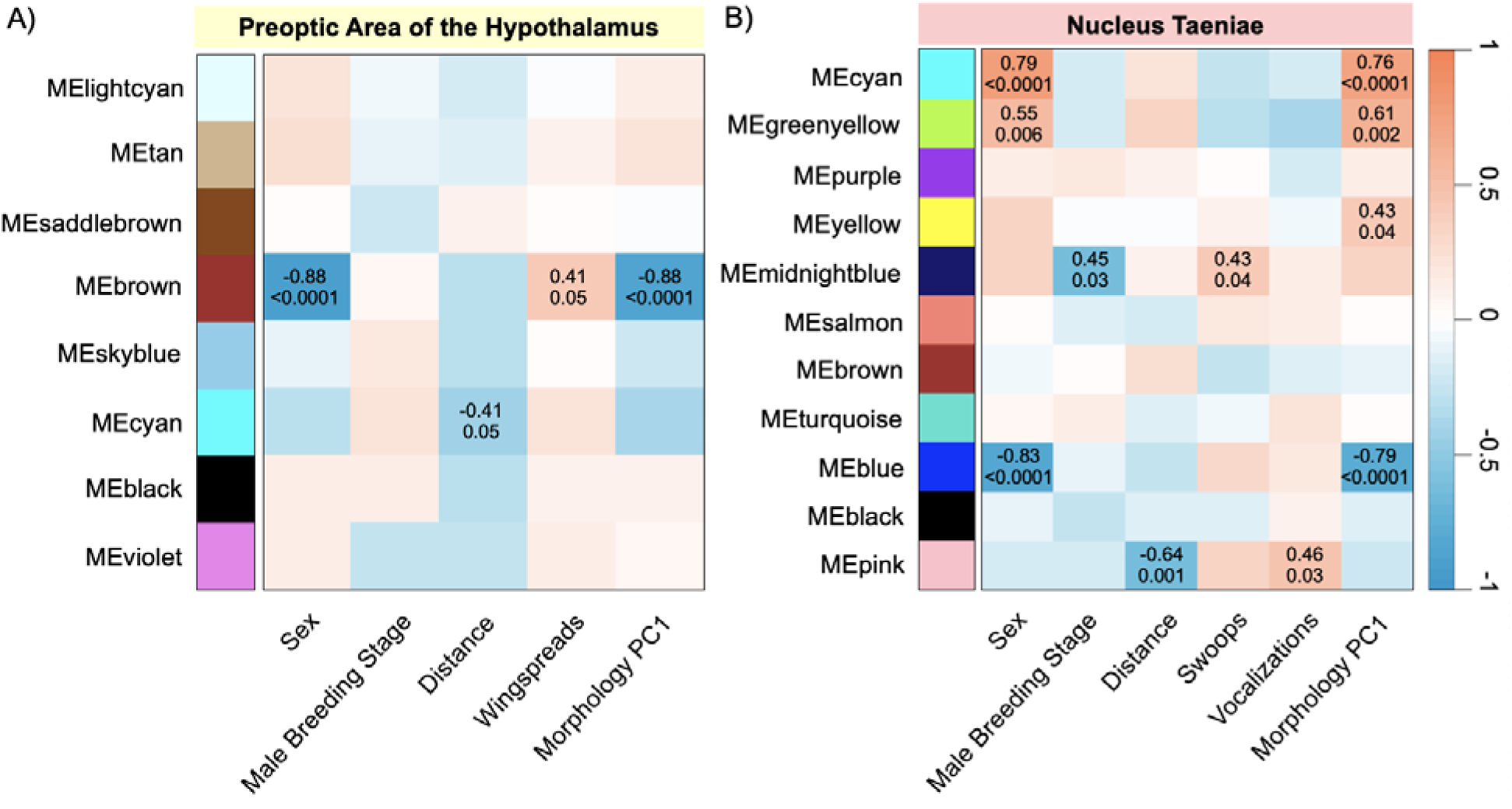
Module-trait relationships in the northern jacana were identified using bicor tests in WGCNA for the a) preoptic area of the hypothalamus and b) the nucleus taeniae. Associations are shown for group (females, courting males, parenting males), sex (males as reference), behavioral responses to resident-intruder assay (distance to decoy, wingspreads, swoops, vocalizations), and size morphology PCA (body mass, tarsus, and wing spur length). Gene modules on the y axis are assigned a random color. The colors on the heatmap are determined by the magnitude and direction of the correlation coefficient. Sample sizes for females (n = 12), courting males (n = 5), and parenting males (n = 7) are combined for all module-trait correlations. Each significant module-trait has the correlation coefficient and the p-value shown. Additional non-significant module-trait relationships are shown in Figures S5, S6.

### Tissue specificity explains most variation in expression

We next examined tissue specificity of gene expression in one representative individual per group. For all samples, most of the 14,588 genes were expressed in all tissues (female 80.8%, courting male 76.7%, parenting male 79.3%; Figure S15). The next largest set for all samples included non-blood tissues, which shared additional genes (female 5.4%, courting male 7.8%, parenting male 5.6%). However, when we normalized expression data from all three individuals, we found that many of these genes were expressed unevenly across tissues (32% of genes with Tau > 0.8; Figure S16), showing strong tissue specificity. Z-linked (29% of genes with Tau > 0.8) and autosomal (32% of genes with Tau > 0.8) genes had similar ratios of uneven expression.

For Z-linked genes, most of the 534 genes were expressed in all tissues, including blood, whole brain, and gonad (56.3%; Figure S17). The next largest set within Z-linked genes included non-blood tissues (19.7%) similar to the full gene dataset. Most of the Z-linked genes that were differentially expressed in the preoptic area of the hypothalamus or nucleus taeniae were also expressed in all tissues.

## DISCUSSION

Investigating sex differences in gene expression in socially polyandrous (i.e. ‘sex-role reversed) species can help illuminate the physiological mechanisms underlying female competition and male parental care. Prior studies of social polyandry identified the potential for increased steroid sensitivity in the brains of females (Voigt and Goymann 2007; Voigt 2016), as well as a sex-independent, conserved set of genes associated with aggression across species (Schumer et al. 2011). Here we extend prior analyses to the northern jacana, a textbook example of social polyandry, by comparing territorial females to males in courting and parenting breeding stages.

### Sex-biased gene expression found on both Z chromosomes and autosomes

We uncovered sex differences in hundreds of genes in the preoptic area of the hypothalamus and the nucleus taeniae (3-4% of brain transcriptome). This proportion of sex-biased genes is similar to socially polyandrous pipefish (Johnson et al. 2024; Pappert et al. 2024) as well as other taxa with diverse mating systems (Santos et al. 2008; Pointer et al. 2013; Whittle et al. 2021).

In the socially-relevant brain regions we examined, most sex-biased genes (83-85%) had higher expression in males. When partitioning Z- and W-linked genes from autosomal genes, we found that most (60-73%) male-biased genes were on the Z chromosome, and most Z-linked genes were male-biased (56-63%). Birds have a ZW-ZZ sex chromosome system; females are the heterogametic sex (ZW), whereas males are the homogametic sex (ZZ).

Dosage compensation on the Z chromosome is incomplete in birds (Itoh 2007; Mank 2013). Thus, male-biased gene expression was likely influenced by the dosage of Z-linked genes. However, a considerable proportion of sex-biased genes were autosomal (40-50%), suggesting that sex differences in gene expression are not simply an artifact of sex chromosomes. Sex differences in brain gene expression persisted regardless of whether males were in courting or parenting breeding stages. We expected females and courting males to have more similar transcriptomic profiles, due to their shared competitive state, but this contrast had a higher proportion of autosomal DEGs in the preoptic area of the hypothalamus. An alternative explanation is that behaviors and molecular mechanisms during courtship are sex-specific (Mank et al. 2013; Ishii et al. 2020), and therefore result in stronger expression differences between females and males. Furthermore, we measured constitutive gene expression, and it is possible that different transcriptomic patterns would emerge in response to social stimulation (Bukhari et al. 2019). Our microdissection approach focused on small, socially-relevant brain regions using bulk RNAseq, and, future work should also consider sex differences in cell type abundance within these tissues, using single-cell transcriptomic sequencing (Darolti and Mank 2023).

We are only beginning to understand the relative roles of Z-linked vs. autosomal genes in shaping sex differences in birds. Several recent studies have associated phenotypes under sex-specific selection with sequence-level variation on Z and W chromosomes (Dunn et al. 2024; Merondun et al. 2024), though in general, the evidence that sexually dimorphic phenotypes are more often encoded by Z-linked genes is limited (Dean and Mank 2014). New proteomic evidence suggests that incomplete dosage compensation of chicken sex chromosomes may be balanced by post-transcriptional changes (Lister et al. 2024), and it will be interesting to connect sex variation in transcriptome to the proteome in more species. Recent work has found that polyandry is associated with stronger selection on Z-linked loci, resulting in faster sequence evolution, known as the faster-Z effect (Wanders et al. 2024), and future work could examine whether sex-biased genes have higher rates of evolution in polyandrous species (Mank et al. 2013). Comparisons of our results with other socially polyandrous species are challenging given the impact of Z-linked genes in birds. However, in cichlid fishes, a microarray study found that the female-dominant *J. marlieri* had more female-biased expression, whereas the male-dominant *J. transcriptus* had more male-biased expression (Schumer et al. 2011).

Studies of socially polyandrous pipefish, which do not have sex determining chromosomes (Dubin et al. 2025), have found that gene expression is more evenly balanced between males and females in the brain and whole-body (Beal et al. 2018; Pappert et al. 2024), and potentially sex-biased in some species (Johnson et al. 2024), depending on analysis methods. Tissue differences may also influence sex-biased patterns of expression (Tosto et al. 2023; Dubin et al. 2025). Similarly, our exploratory analyses of brain, blood and gonad found that many genes were unevenly expressed across tissues, though most Z-linked and autosomal genes were expressed in all tissue types, with similar ratios of uneven expression. Across taxa, the lack of consistent sex differences in gene expression suggest that social polyandry is not a simple reversal in the direction of sex-biased gene expression.

### Male breeding stage is associated with few changes in autosomal gene expression

Though our study was designed to compare gene expression between females and males, these males were in two different breeding stages, allowing us to compare gene expression in courtship versus parenting, albeit with smaller sample sizes. Relative to sex differences, few genes differed in expression between male courting and parenting stages. There were 19 differentially expressed autosomal genes in the preoptic area of the hypothalamus, and none in the nucleus taeniae. Our results are similar to a study in male uniparental poison frogs (*Dendrobates tinctorus),* which found few differentially expressed genes between stages of egg care versus no parental care (Fischer and O’Connell 2020). However, our results contrast with other studies finding hundreds of neurogenomic changes associated with the transition into parental care in male sticklebacks (Bukhari et al. 2019) and female tree swallows (Bentz et al. 2019). We observed males when parental care was already established, and perhaps may have found more transcriptomic changes in the immediate transition between courtship and parenting, particularly if neurotranscriptional changes are most pronounced during a social transition like they are in some other settings (Maruska et al. 2013). Alternatively, male jacanas may remain in a ‘ready-to-parent’ state as they cycle between courtship and parenting throughout the breeding season. This opens new questions on the moment-to-moment experiences that shift their physiology and behavior so dramatically across breeding stages, and the extent to which more rapid neuromodulatory regulation shapes this transition.

Many of the genes that differed in expression between male breeding stages relate to hormonal actions associated with reproduction. Courting males had higher expression of the opioid neuropeptide precursor prodynorphin (*PDYN*), which has previously been associated with nesting and singing behaviors in male European starlings *Sturnus vulgaris* (Riters et al. 2017).

Three genes related to the thyroid and insulin-like growth factor (*IGF*) pathways, Iodotyrosine Deiodinase (IYD), *IGF* Binding Protein 3 (*IGFBP3*) and Pappalysin 2 (*PAPPA2*), were more highly expressed in parenting males than in courting males. The thyroid and IGF pathways are associated with seasonal reproduction in birds (Sharp 2005; Verhagen et al. 2019). The stage-specific transcriptomic changes we did observe were only in the preoptic area of the hypothalamus, which has been linked to parental care and competitive behaviors across many species of vertebrates. In contrast, the nucleus taeniae may be more related to social and competitive valence (Goodson and Wang 2006) in ways that are relevant regardless of stage. Future work should examine additional nodes of the vertebrate social behavior network and how they interact, particularly because there is potential for even subtle differences in one node to be amplified as behavioral circuits are activated across the entire brain (Goodson 2005).

Emerging data suggest that these multi-region networks operate differently for behaviorally distinct individuals (Bolton et al. 2025), suggesting they also may vary within an individual, too.

### Gene expression related to hormone signaling may reflect competitive and parental behaviors

Studies of socially polyandrous species, including black coucals and barred buttonquails, found similar patterns in which *AR* expression was higher in females than males, though in the nucleus taeniae instead of the preoptic area (Voigt and Goymann 2007; Voigt 2016). A previous study of testosterone levels in these same individuals found that females have similar levels of testosterone to parenting males, and that testosterone levels were highest in courting males (Lipshutz and Rosvall 2020b). Though female jacanas have low levels of circulating testosterone, higher androgen sensitivity through receptor expression could promote female competition, and future work could experimentally test this relationship, as well as the role of other androgens including androstenedione. Androgen-mediated phenotypes are regulated not only by testosterone levels, but also by mechanisms of steroid metabolism and sensitivity (Fuxjager and Schuppe 2018; Loveland et al. 2025), including the binding of testosterone to androgen receptors across tissue types.

Aromatase can locally convert testosterone to estradiol in the brain (Schmidt et al. 2008), and this mechanism has been shown to regulate aggression and sexual behavior in many species (Trainor et al. 2006). In the preoptic area of the hypothalamus, aromatase (*CYP19A1*) expression was marginally higher in courting males than parenting males, but did not differ between sexes. Estrogen receptor 1 (*ESR1)* was higher in females in the preoptic area of the hypothalamus. Androgen (*AR*) and estrogen (*ESR1, ESR2*) receptor expression did not differ between male breeding stages for either brain region. Altogether, this suggests that differences in androgen production and potentially conversion may shape stage-specific differences in courtship and parental care in male jacanas, in contrast to androgen reception.

We found that prolactin receptor (*PRLR*) expression was male-biased in both brain regions, and *PRLR* expression was marginally higher in parenting males than in courting males in the preoptic area of the hypothalamus, a region particularly important in regulating parental care (Smiley 2019). Prolactin signaling promotes parental behavior in both females and males of other avian species (Angelier and Chastel 2009). Previous studies of socially polyandrous birds found higher prolactin levels in males than females (Oring et al. 1986; Gratto-Trevor et al. 1990). Surprisingly, prolactin-releasing hormone (*PRLH*), which is one mechanism by which prolactin levels can increase (Spuch et al. 2007), was more highly expressed in courting males compared to parenting males. PRLR is also Z-linked, and higher expression has been found in males than females in another bird species (Farrar et al. 2022). In northern jacanas, higher expression of *PRLR* may act as a mechanism for paternal care, on top of the Z-linked effect.

Our results suggest that sex differences in behavior may result from complex interactions between endocrine pathways and gene expression, and highlight how gonadal sex versus parental roles may shape transcriptomic variation between females and males.

### Competitive traits are correlated with gene modules

In addition to comparing sex differences in gene expression, we also used co-expression network analyses to connect variation in gene expression with competitive traits. Sexually dimorphic gene expression was apparent in several gene modules, and we infer that most of these module-trait correlations were driven by confounded sex differences in both gene expression and phenotypic traits. Across brain regions, two male-biased modules overlapped significantly in the makeup of genes and were primarily made up of Z-linked genes. However, there were several gene modules associated with aggression but not with sex differences, and these may provide insights into variation between gene expression and competitive behaviors. In the preoptic area of the hypothalamus, the coexpression of genes in the cyan module correlated positively with aggression (closer distance to decoy) and was enriched for central nervous system myelination, a process associated with social stress and aggression in mice (Bonnefil et al. 2019). In the nucleus taeniae, genes in the pink module correlated positively with aggression, including closer distance to decoy and more vocalizations. The genes in this module were enriched for biological processes including regulation of cilium beat frequency and sperm axoneme assembly, though in the brain, this may instead reflect microtubule-associated neurobiological processes like axonal or dendritic growth.

### Disentangling molecular mechanisms of ‘sex roles’ from sex

Socially polyandrous species provide a unique opportunity to identify the extent to which physiological differences between females and males are shaped by sex roles. In the northern jacana, we found patterns of sex-biased gene expression explained by both Z-linked and autosomal genes. These patterns of sex-specific gene expression are similar to other bird species with ‘traditional’ sex roles of male competition and female parental care (Arnold and Itoh 2011). Contrary to expectations that female and courting males would have similar neurogenomic profiles, autosomal gene expression differed most between these groups.

However, sex differences in brain gene expression persisted regardless of male breeding stage, suggesting alternative mechanisms of regulation as males transition from courting to parenting across the breeding season, including hormone signaling (Lipshutz and Rosvall 2020b). We also saw sex-specific expression of genes associated with androgen and prolactin signaling, which may reflect hormonal sensitivity associated with competition and parental care. We conclude that social polyandry (i.e., ‘sex-role reversal’) is not a simple reversal in the direction of sex-biased gene expression in the brain, but rather a result of complex genetic and hormonal interactions. Future work comparing socially polyandrous species with close relatives that have ‘traditional’ sex roles would be instructive for identifying genes involved in the transition to social polyandry.

## Notes

https://github.com/tpatton4/jacananeurogenomics

### Competing Interest Statement

The authors have declared no competing interest.

### Funding

- U.S. National Science Foundation: IOS-1818235, DBI-1907134
- National Institutes of Health: T32HD049336

## Reference

Ah-King, M. 2013. On anisogamy and the evolution of ‘sex roles’. Trends Ecol Evol 28:1–2.

Alonge, M., L. Lebeigle, M. Kirsche, K. Jenike, S. Ou, S. Aganezov, X. Wang, Z. B. Lippman, M. C. Schatz, and S. Soyk. 2022. Automated assembly scaffolding using RagTag elevates a new tomato system for high-throughput genome editing. Genome Biol 23:258.

Angelier, F. and O. Chastel. 2009. Stress, prolactin and parental investment in birds: A review. General and Comparative Endocrinology 163:142–148.

Arnold, A. P. and Y. Itoh. 2011. Factors causing sex differences in birds. Avian Biol Res 4.

Bateman, A. J. 1948. Intra-sexual selection in Drosophila. Heredity 2:349–368.

Beal, A. P., F. D. Martin, and M. C. Hale. 2018. Using RNA-seq to determine patterns of sex-bias in gene expression in the brain of the sex-role reversed Gulf Pipefish (Syngnathus scovelli). Mar Genomics 37:120–127.

Bentz, A. B., D. B. Rusch, A. Buechlein, and K. A. Rosvall. 2019. The neurogenomic transition from territory establishment to parenting in a territorial female songbird. BMC Genomics 20:819.

Bolger, A. M., M. Lohse, and B. Usadel. 2014. Trimmomatic: a flexible trimmer for Illumina sequence data. Bioinformatics 30:2114–2120.

Bolton, P. E., T. B. Ryder, R. Dakin, J. L. Houtz, I. T. Moore, C. N. Balakrishnan, and B. M. Horton. 2025. Neurogenomic landscape associated with status-dependent cooperative behaviour. Mol Ecol 34:e17327.

Bonnefil, V., K. Dietz, M. Amatruda, M. Wentling, A. V. Aubry, J. L. Dupree, G. Temple, H. J. Park, N. S. Burghardt, P. Casaccia, and J. Liu. 2019. Region-specific myelin differences define behavioral consequences of chronic social defeat stress in mice. Elife 8.

Bukhari, S. A., M. C. Saul, N. James, M. K. Bensky, L. R. Stein, R. Trapp, and A. M. Bell. 2019. Neurogenomic insights into paternal care and its relation to territorial aggression. Nat Commun 10:4437.

Burmeister, S. S., E. D. Jarvis, and R. D. Fernald. 2005. Rapid behavioral and genomic responses to social opportunity. PLoS Biol 3:e363.

Cheng, H., G. T. Concepcion, X. Feng, H. Zhang, and H. Li. 2021. Haplotype-resolved de novo assembly using phased assembly graphs with hifiasm. Nat Methods 18:170–175.

Darolti, I. and J. E. Mank. 2023. Sex-biased gene expression at single-cell resolution: cause and consequence of sexual dimorphism. Evol Lett 7:148–156.

Dean, R. and J. E. Mank. 2014. The role of sex chromosomes in sexual dimorphism: discordance between molecular and phenotypic data. J Evol Biol 27:1443–1453.

Dubin, A., J. Parker, A. Bohne, and O. Roth. 2025. Sexual Antagonism and Sex Determination in Three Syngnathid Species Alongside a Male Pregnancy Gradient. Genome Biol Evol 17.

Dunn, P. O., N. D. Sly, C. R. Freeman-Gallant, A. E. Henschen, C. M. Bossu, K. C. Ruegg, P. Minias, and L. A. Whittingham. 2024. Sexually selected differences in warbler plumage are related to a putative inversion on the Z chromosome. Mol Ecol 33:e17525.

Emlen, S. T. and L. W. Oring. 1977. Ecology, sexual selection, and the evolution of mating systems. Science 197:215–223.

Emlen, S. T. and P. H. Wrege. 2004. Size Dimorphism, Intrasexual Competition, and Sexual Selection in Wattled Jacana (Jacana jacana), a sex-role-reversed shorebird in Panama. The Auk 121:391–403.

Farrar, V. S., R. M. Harris, S. H. Austin, B. M. Nava Ultreras, A. M. Booth, F. Angelier, A. S. Lang, T. Feustel, C. Lee, A. Bond, M. D. MacManes, and R. M. Calisi. 2022. Prolactin and prolactin receptor expression in the HPG axis and crop during parental care in both sexes of a biparental bird (Columba livia). Gen Comp Endocrinol 315:113940.

Feldman, R., K. Braun, and F. A. Champagne. 2019. The neural mechanisms and consequences of paternal caregiving. Nat Rev Neurosci 20:205–224.

Fischer, E. K. and L. A. O’Connell. 2020. Hormonal and neural correlates of care in active versus observing poison frog parents. Horm Behav 120:104696.

Fuxjager, M. J. and E. R. Schuppe. 2018. Androgenic signaling systems and their role in behavioral evolution. The Journal of Steroid Biochemistry and Molecular Biology 184:47–56.

Goodson, J. L. 2005. The vertebrate social behavior network: evolutionary themes and variations. Horm Behav 48:11–22.

Goodson, J. L. and Y. Wang. 2006. Valence-sensitive neurons exhibit divergent functional profiles in gregarious and asocial species. Proc Natl Acad Sci USA 103:17013–17017.

Grath, S. and J. Parsch. 2016. Sex-Biased Gene Expression. Annu Rev Genet 50:29–44.

Gratto-Trevor, C. L., L. W. Fivizzani Aj Fau - Oring, F. Oring Lw Fau - Cooke, and F. Cooke. 1990. Seasonal changes in gonadal steroids of a monogamous versus a polyandrous shorebird. Gen Comp Endocrinol 80:407–418.

Horton, B. M., W. H. Hudson, E. A. Ortlund, S. Shirk, J. W. Thomas, E. R. Young, W. M. Zinzow-Kramer, and D. L. Maney. 2014. Estrogen receptor alpha polymorphism in a species with alternative behavioral phenotypes. Proc Natl Acad Sci USA 111:1443–1448.

Horton, B. M., T. B. Ryder, I. T. Moore, and C. N. Balakrishnan. 2020. Gene expression in the social behavior network of the wire-tailed manakin (Pipra filicauda) brain. Genes Brain Behav 19:e12560.

Houtz, J. L., K. A. Acosta, M. Berlow, and S. E. Lipshutz. 2025. Reproductive microbial diversity is associated with competitive phenotypes in socially polyandrous jacanas. BioRxiv.

Hudson, N. J., A. Reverter, and B. P. Dalrymple. 2009. A Differential Wiring Analysis of Expression Data Correctly Identifies the Gene Containing the Causal Mutation. PLOS Computational Biology 5:e1000382.

Ishii, K., M. Wohl, A. DeSouza, and K. Asahina. 2020. Sex-determining genes distinctly regulate courtship capability and target preference via sexually dimorphic neurons. Elife 9.

Itoh, Y., Melamed, E., Yang, X. et al. . 2007. Dosage compensation is less effective in birds than in mammals. Journal of Biology 6.

Janicke, T., I. K. HalJderer, M. J. Lajeunesse, and N. Anthes. 2016. Darwinian sex roles confirmed across the animal kingdom. Science Advances 2.

Jenni, D. A. and G. Collier. 1972. Polyandry in the American Jaçana (Jacana spinosa). The Auk 89:743–765.

Johnson, B. D., E. Rose, and A. G. Jones. 2024. Sensitivity of transcriptomics: Different samples and methodology alter conclusions in Gulf pipefish (Syngnathus scovelli). J Hered.

Kim, D., J. M. Paggi, C. Park, C. Bennett, and S. L. Salzberg. 2019. Graph-based genome alignment and genotyping with HISAT2 and HISAT-genotype. Nat Biotechnol 37:907–915.

Kohl, J., A. E. Autry, and C. Dulac. 2017. The neurobiology of parenting: A neural circuit perspective. Bioessays 39:1–11.

Kretschmer, R., M. S. Souza, S. A. Barcellos, T. M. Degrandi, J. C. Pereira, P. C. M. O’Brien, M. A. Ferguson-Smith, R. J. Gunski, A. D. V. Garnero, E. H. C. Oliveira, and T. R. O. Freitas. 2020. Novel insights into chromosome evolution of Charadriiformes: extensive genomic reshuffling in the wattled jacana (Jacana jacana, Charadriiformes, Jacanidae). Genet Mol Biol 43:e20190236.

Kuenzel, W. J. and C. D. Golden. 2006. Distribution and change in number of gonadotropin-releasing hormone-1 neurons following activation of the photoneuroendocrine system in the chick, Gallus gallus. Cell Tissue Res 325:501–512.

Langfelder, P. and S. Horvath. 2008. WGCNA: an R package for weighted correlation network analysis. BMC Bioinformatics 9:559.

Langmead, B. and S. L. Salzberg. 2012. Fast gapped-read alignment with Bowtie 2. Nat Methods 9:357–359.

Li, H. 2018. Minimap2: pairwise alignment for nucleotide sequences. Bioinformatics 34:3094–3100.

Li, H., B. Handsaker, A. Wysoker, T. Fennell, J. Ruan, N. Homer, G. Marth, G. Abecasis, R. Durbin, and S. Genome Project Data Processing. 2009. The Sequence Alignment/Map format and SAMtools. Bioinformatics 25:2078–2079.

Liao, Y., G. K. Smyth, and W. Shi. 2019. The R package Rsubread is easier, faster, cheaper and better for alignment and quantification of RNA sequencing reads. Nucleic Acids Res 47:e47.

Lipshutz, S. E. 2017. Divergent competitive phenotypes between females of two sex-role-reversed species. Behavioral Ecology and Sociobiology 71.

Lipshutz, S. E. and K. A. Rosvall. 2020a. Neuroendocrinology of Sex-Role Reversal. Integr Comp Biol 60:692–702.

Lipshutz, S. E. and K. A. Rosvall. 2020b. Testosterone secretion varies in a sex- and stage-specific manner: Insights on the regulation of competitive traits from a sex-role reversed species. Gen Comp Endocrinol 292:113444.

Lipshutz, S. E., S. J. Torneo, and K. A. Rosvall. 2023. How Female-Female Competition Affects Male-Male Competition: Insights into Postcopulatory Sexual Selection from Socially Polyandrous Species. Am Nat 201:460–471.

Lister, N. C., A. M. Milton, H. R. Patel, S. A. Waters, B. J. Hanrahan, K. L. McIntyre, A. M. Livernois, W. B. Horspool, L. K. Wee, A. R. Ringel, S. Mundlos, M. I. Robson, L. Shearwin-Whyatt, F. Grutzner, J. A. M. Graves, A. Ruiz-Herrera, and P. D. Waters. 2024. Incomplete transcriptional dosage compensation of chicken and platypus sex chromosomes is balanced by post-transcriptional compensation. Proc Natl Acad Sci U S A 121:e2322360121.

Loveland, J. L., A. Zemella, V. M. Jovanović, G. Möller, C. P. Sager, B. Bastos, K. A. Dyar, L. Fusani, M. Gahr, L. M. Giraldo-Deck, W. Goymann, D. B. Lank, J. Tokarz, K. Nowick, and C. Küpper. 2025. A single gene orchestrates androgen variation underlying male mating morphs in ruffs. Science 387:406–412.

Luna, L. W. and S. E. Lipshutz. Genomic signals of reproductive character displacement in a bird hybrid zone, Unpublished data.

Mank, J. E. 2013. Sex chromosome dosage compensation: definitely not for everyone. Trends Genet 29:677–683.

Mank, J. E., N. Wedell, and D. J. Hosken. 2013. Polyandry and sex-specific gene expression. Philos Trans R Soc Lond B Biol Sci 368:20120047.

Maruska, K. P., L. Becker, A. Neboori, and R. D. Fernald. 2013. Social descent with territory loss causes rapid behavioral, endocrine and transcriptional changes in the brain. J Exp Biol 216:3656–3666.

Merondun, J., C. I. Marques, P. Andrade, S. Meshcheryagina, I. Galván, S. Afonso, J. M. Alves, P. M. Araújo, G. Bachurin, J. Balacco, M. Bán, O. Fedrigo, G. Formenti, F. Fossøy, A. Fülöp, M. Golovatin, S. Granja, C. Hewson, M. Honza, K. Howe, G. Larson, A. Marton, C. Moskát, J. Mountcastle, P. Procházka, Y. Red’kin, Y. Sims, M. Šulc, A. Tracey, J. M. D. Wood, E. D. Jarvis, M. E. Hauber, M. Carneiro, and J. B. W. Wolf. 2024. Evolution and genetic architecture of sex-limited polymorphism in cuckoos. Science Advances 10:eadl5255.

Mi, H., A. Muruganujan, X. Huang, D. Ebert, C. Mills, X. Guo, and P. D. Thomas. 2019. Protocol Update for large-scale genome and gene function analysis with the PANTHER classification system (v.14.0). Nat Protoc 14:703–721.

Muck, C. and W. Goymann. 2011. Throat patch size and darkness covaries with testosterone in females of a sex-role reversed species. Behavioral Ecology 22:1312–1319.

Munley, K. M., K. L. Wade, and D. S. Pradhan. 2022. Uncovering the seasonal brain: Liquid chromatography-tandem mass spectrometry (LC-MS/MS) as a biochemical approach for studying seasonal social behaviors. Horm Behav 142:105161.

Nursyifa, C., A. Bruniche-Olsen, G. Garcia-Erill, R. Heller, and A. Albrechtsen. 2022. Joint identification of sex and sex-linked scaffolds in non-model organisms using low depth sequencing data. Mol Ecol Resour 22:458–467.

Oring, L. W., A. J. Fivizzani, M. E. el Halawani, and A. Goldsmith. 1986. Seasonal changes in prolactin and luteinizing hormone in the polyandrous spotted sandpiper, Actitis macularia. General and Comparative Endocrinology 62:394–403.

Pappert, F. A., D. Kolbe, A. Dubin, and O. Roth. 2024. The effect of parental age on the quantity and quality of offspring in Syngnathus typhle, a species with male pregnancy. Evol Appl 17:e13755.

Pointer, M. A., P. W. Harrison, A. E. Wright, and J. E. Mank. 2013. Masculinization of gene expression is associated with exaggeration of male sexual dimorphism. PLoS Genet 9:e1003697.

Puelles, L. 2007. The Chick Brain in Stereotaxic Coordinates : An Atlas Featuring Neuromeric Subdivisions and Mammalian Homologies. Academic Press, Amsterdam.

Riters, L. V., M. A. Cordes, and S. A. Stevenson. 2017. Prodynorphin and kappa opioid receptor mRNA expression in the brain relates to social status and behavior in male European starlings. Behav Brain Res 320:37–47.

Rosvall, K. A., A. B. Bentz, and E. M. George. 2020. How research on female vertebrates contributes to an expanded challenge hypothesis. Horm Behav 123:104565.

Rosvall, K. A., C. M. Bergeon Burns, J. Barske, J. L. Goodson, B. A. Schlinger, D. R. Sengelaub, and E. D. Ketterson. 2012. Neural sensitivity to sex steroids predicts individual differences in aggression: implications for behavioural evolution. Proc Biol Sci 279:3547–3555.

Santos, E. M., P. Kille, V. L. Workman, G. C. Paull, and C. R. Tyler. 2008. Sexually dimorphic gene expression in the brains of mature zebrafish. Comp Biochem Physiol A Mol Integr Physiol 149:314–324.

Schmidt, K. L., D. S. Pradhan, A. H. Shah, T. D. Charlier, E. H. Chin, and K. K. Soma. 2008. Neurosteroids, immunosteroids, and the Balkanization of endocrinology. Gen Comp Endocrinol 157:266–274.

Schumer, M., K. Krishnakant, and S. C. Renn. 2011. Comparative gene expression profiles for highly similar aggressive phenotypes in male and female cichlid fishes (Julidochromis). J Exp Biol 214:3269–3278.

Sharp, P. J. 2005. Photoperiodic regulation of seasonal breeding in birds. Ann N Y Acad Sci 1040:189–199.

Smiley, K. O. 2019. Prolactin and avian parental care: New insights and unanswered questions. Horm Behav 111:114–130.

Smiley, K. O., K. M. Munley, K. Aghi, S. E. Lipshutz, T. M. Patton, D. S. Pradhan, T. K. Solomon-Lane, and S. E. D. Sun. 2024. Sex diversity in the 21st century: Concepts, frameworks, and approaches for the future of neuroendocrinology. Horm Behav 157:105445.

Spuch, C., Y. Diz-Chaves, D. Perez-Tilve, M. Alvarez-Crespo, and F. Mallo. 2007. Prolactin-releasing Peptide (PrRP) increases prolactin responses to TRH in vitro and in vivo. Endocrine 31:119–124.

Stelzer, G., N. Rosen, I. Plaschkes, S. Zimmerman, M. Twik, S. Fishilevich, T. I. Stein, R. Nudel, I. Lieder, Y. Mazor, S. Kaplan, D. Dahary, D. Warshawsky, Y. Guan-Golan, A. Kohn, N. Rappaport, M. Safran, and D. Lancet. 2016. The GeneCards Suite: From Gene Data Mining to Disease Genome Sequence Analyses. Current Protocols in Bioinformatics 54:1.30.31-31.30.33.

Tosto, N. M., E. R. Beasley, B. B. M. Wong, J. E. Mank, and S. P. Flanagan. 2023. The roles of sexual selection and sexual conflict in shaping patterns of genome and transcriptome variation. Nat Ecol Evol 7:981–993.

Trainor, B. C., H. H. Kyomen, and C. A. Marler. 2006. Estrogenic encounters: how interactions between aromatase and the environment modulate aggression. Front Neuroendocrinol 27:170–179.

Trivers, R. L. 1972. Parental investment and sexual selection. Sexual selection and the descent of man. Aldine, Chicago.

Verhagen, I., V. N. Laine, A. C. Mateman, A. Pijl, R. de Wit, B. van Lith, W. Kamphuis, H. M. Viitaniemi, T. D. Williams, S. P. Caro, S. L. Meddle, P. Gienapp, K. van Oers, and M. E. Visser. 2019. Fine-tuning of seasonal timing of breeding is regulated downstream in the underlying neuro-endocrine system in a small songbird. J Exp Biol 222.

Voigt, C. 2016. Neuroendocrine correlates of sex-role reversal in barred buttonquails. Proc Biol Sci 283.

Voigt, C. and W. Goymann. 2007. Sex-role reversal is reflected in the brain of African black coucals (Centropus grillii). Dev Neurobiol 67:1560–1573.

Wanders, K., G. Chen, S. Feng, T. Szekely, and A. O. Urrutia. 2024. Role-reversed polyandry is associated with faster fast-Z in shorebirds. Proc Biol Sci 291:20240397.

Whittle, C. A., A. Kulkarni, and C. G. Extavour. 2021. Evolutionary dynamics of sex-biased genes expressed in cricket brains and gonads. J Evol Biol 34:1188–1211.

Wingfield, J. C., R. E. Hegner, J. Dufty, Alfred M., and G. F. Ball. 1990. The “Challenge Hypothesis”: Theoretical Implications for Patterns of Testosterone Secretion, Mating Systems, and Breeding Strategies. The American Naturalist 136:829–846.

Zilkha, N., N. Scott, and T. Kimchi. 2017. Sexual Dimorphism of Parental Care: From Genes to Behavior. Annu Rev Neurosci 40:273–305.

