## Supplementary material for "Sex and breeding stage differences in neurogenomic profiles reflect hormone signaling in a socially polyandrous shorebird": Figure S1

**Supplemental Figures**

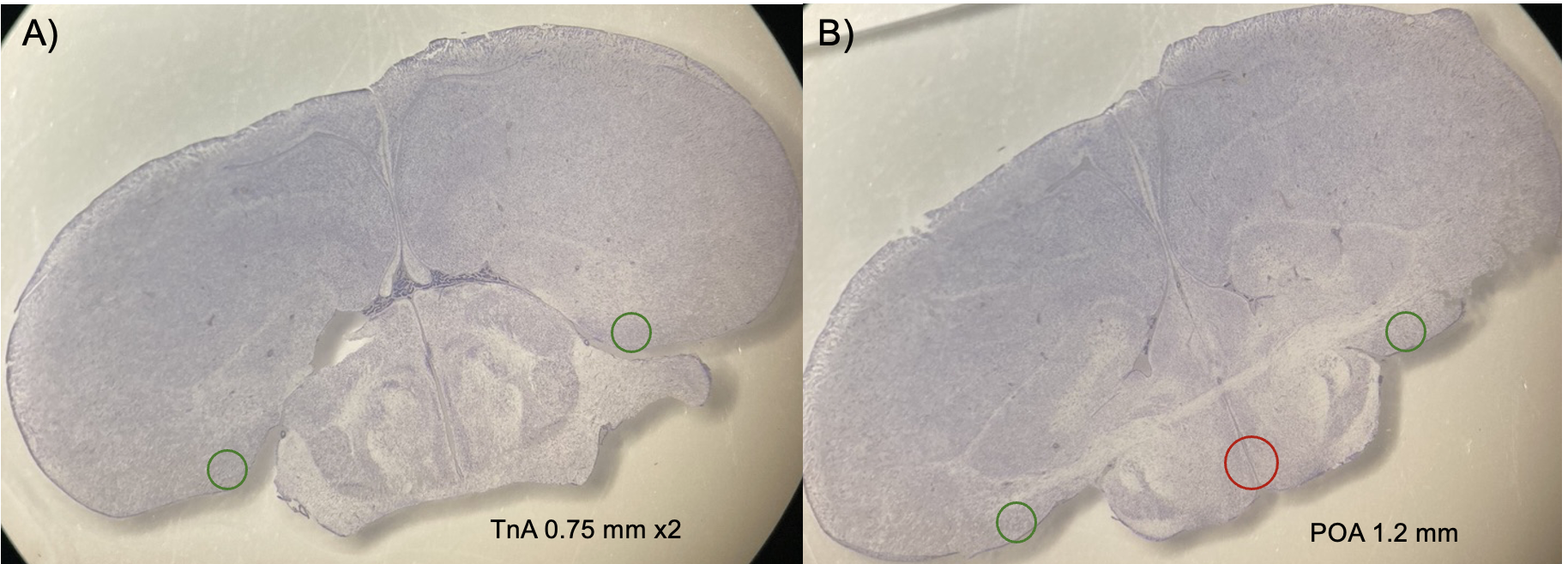

**Figure S1.** Stained cresyl violet sections of a jacana brain with micropunches indicating dissection for the A) nucleus taeniae (TnA), where both hemispheres were punched at 0.75mm or 0.5mm and B) preoptic area of the hypothalamus (POA), punched at 1.2mm, with bilateral TnA in the same section. Coronal sections were conducted from caudal to rostral. Jacana brain is ~16.mm wide and 9.5mm tall.

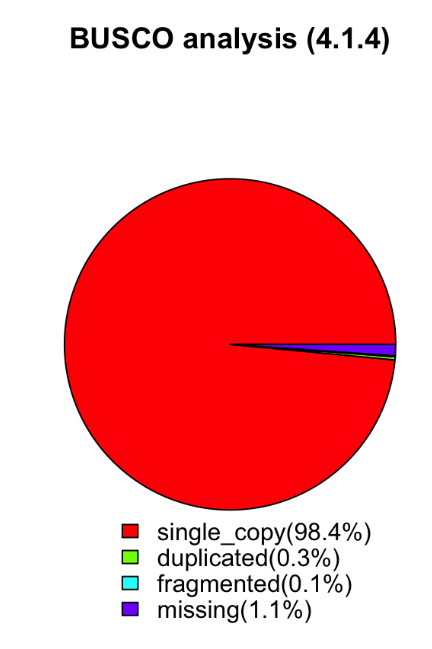

**Figure S2.** BUSCO assessment of completeness of the *Jacana spinosa* HiFi genome showing 98.8% (8,234) of the conserved genes in the data set are complete, with 98.4% (8,207) present as single-copy and 0.3% (27) as duplicated. Fragmented and missing genes account for 0.1% (11) and 1.1% (93), respectively. Additionally, 6.9% (570) of the analyzed genes contain internal stop codons.

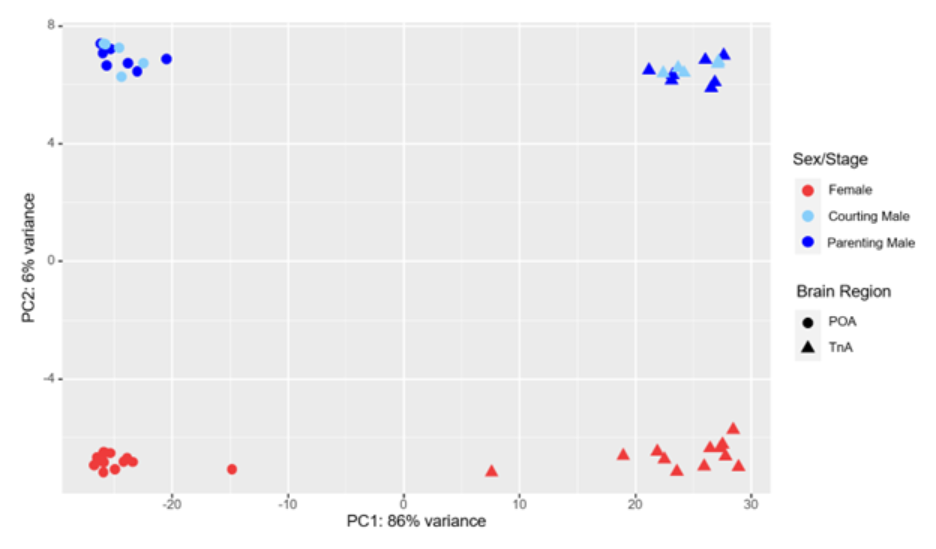

**Figure S3.** Principal component analysis (PCA) plot of differential expression data across all samples, including females (red), courting males (light blue), and parenting males (dark blue).

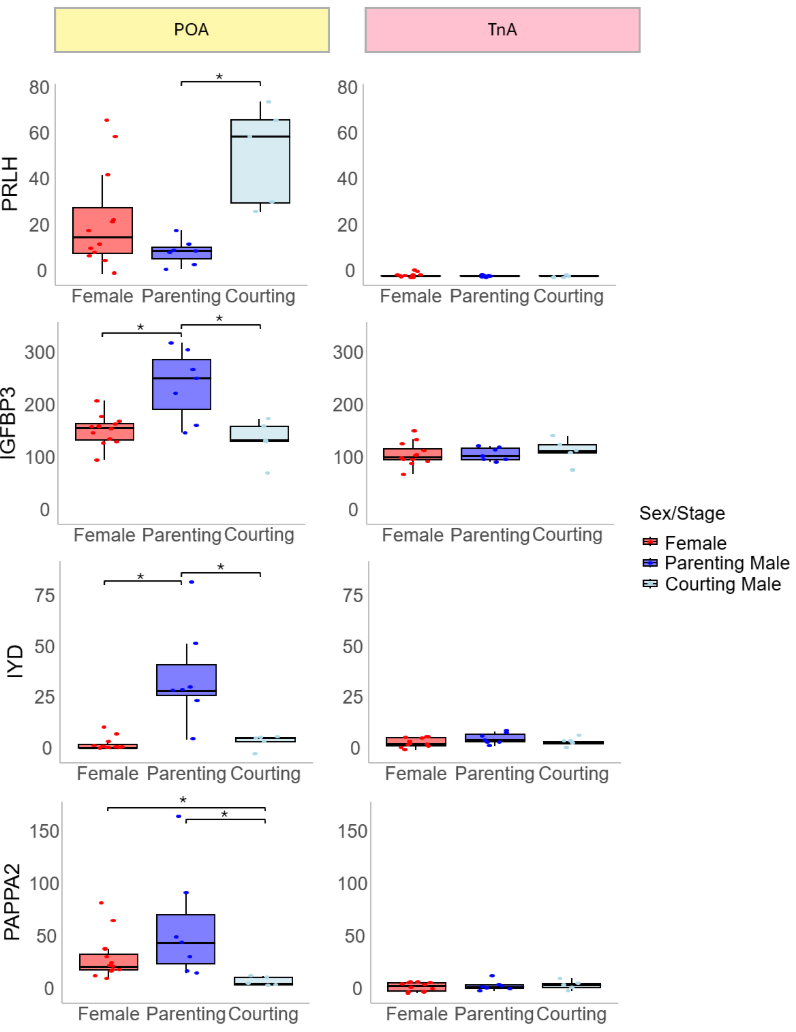

**Figure S4.** Boxplots of genes significantly differentially expressed between parenting and courting males in the preoptic area of the hypothalamus (POA). Boxplots on the right show expression in the nucleus taeniae (TnA), though these genes were not differentially expressed in this brain region. All genes are located on autosomes. *=p<0.05 after false discovery rate.

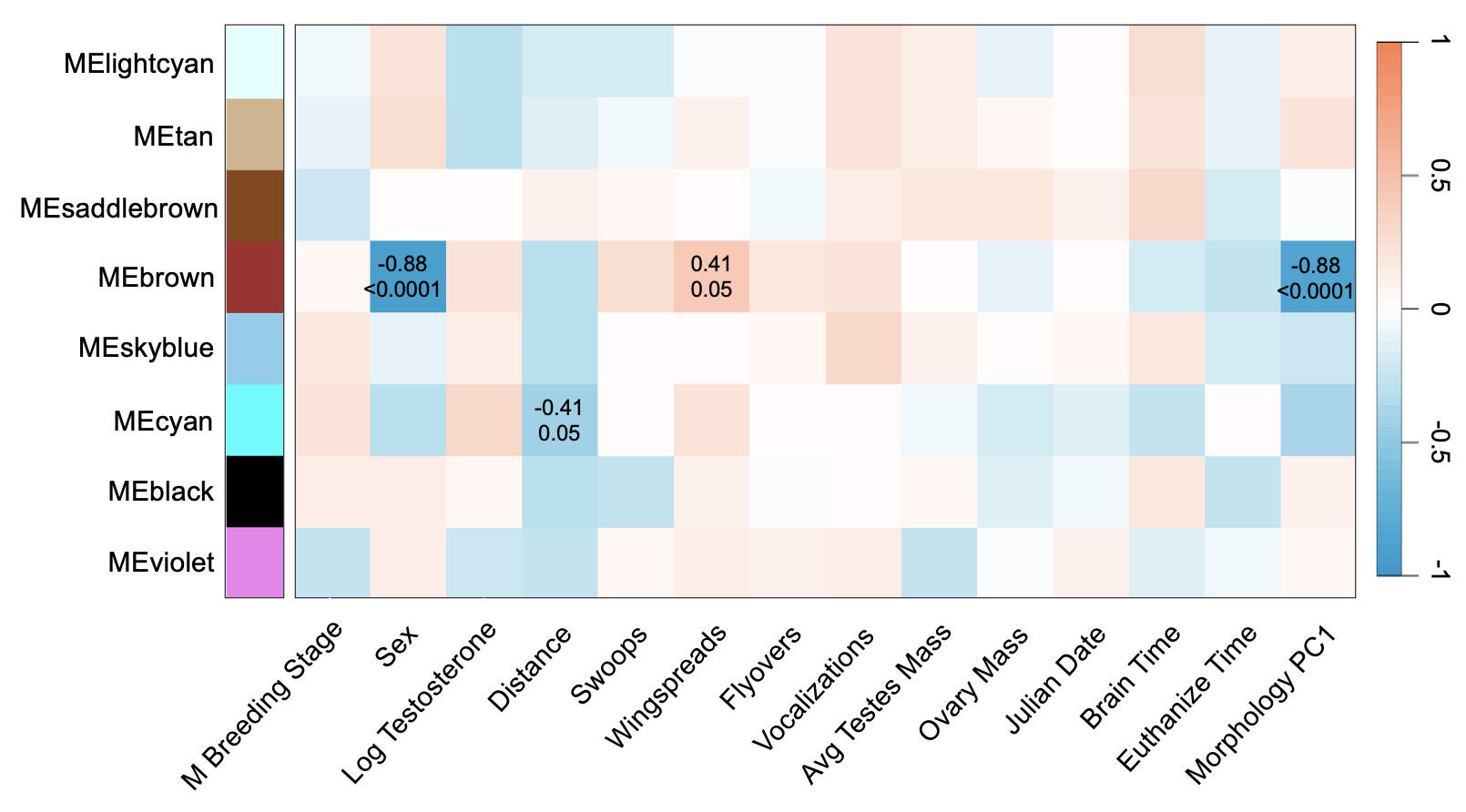

**Figure S5**. Extended module-trait relationships in the northern jacana preoptic area of the hypothalamus were identified using bicor tests in WGCNA. Associations are shown for male breeding stage (courting males, parenting males), sex (females as reference), log testosterone, behavioral responses to resident-intruder assay (distance to decoy, swoops, wing spread, flyovers, vocalizations), gonad mass (testes, ovary), julian date, brain dissection time, time from assay start to euthanasia, and morphological traits size (body mass, tarsus length, and wing spur). Correlation coefficients are shown above with p-values below.

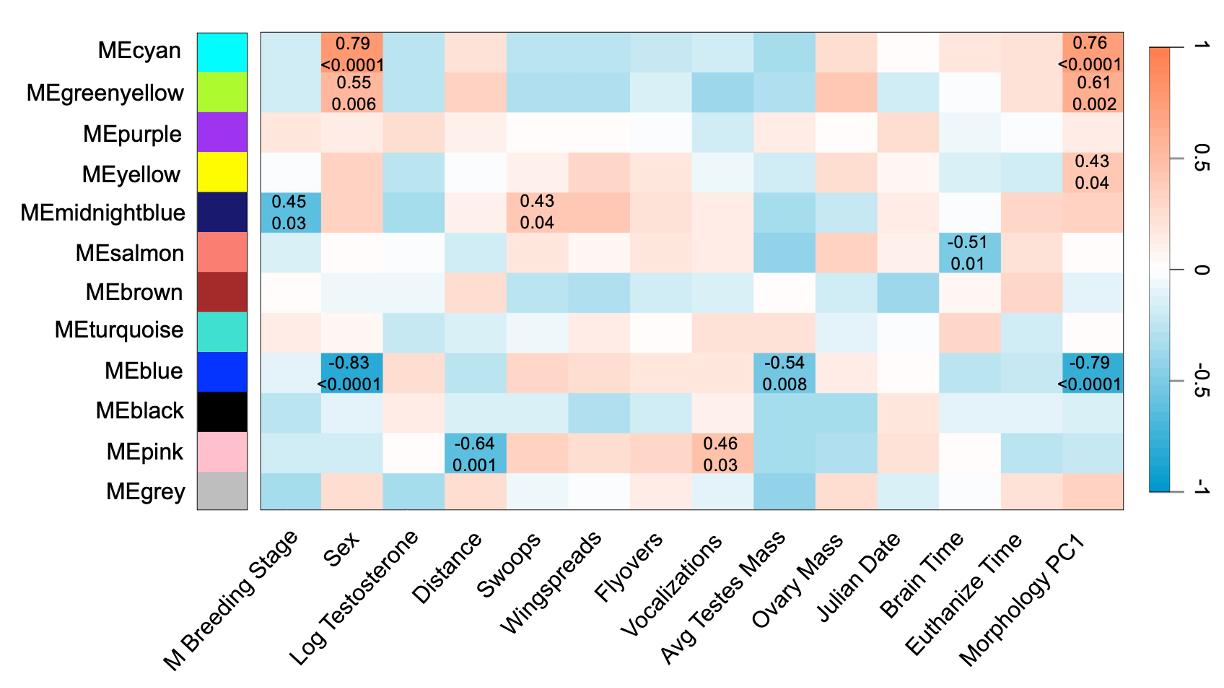

**Figure S6**. Extended module-trait relationships in the northern jacana nucleus taeniae were identified using bicor tests in WGCNA. Associations are shown for male breeding stage (courting males, parenting males), sex (females as reference), log testosterone, behavioral responses to resident-intruder assay (distance to decoy, swoops, wing spread, flyovers, vocalizations), gonad mass (testes, ovary), julian date, brain dissection time, time from assay start to euthanasia, and morphological traits size (body mass, tarsus length, and wing spur). Correlation coefficients are shown above with p-values below.

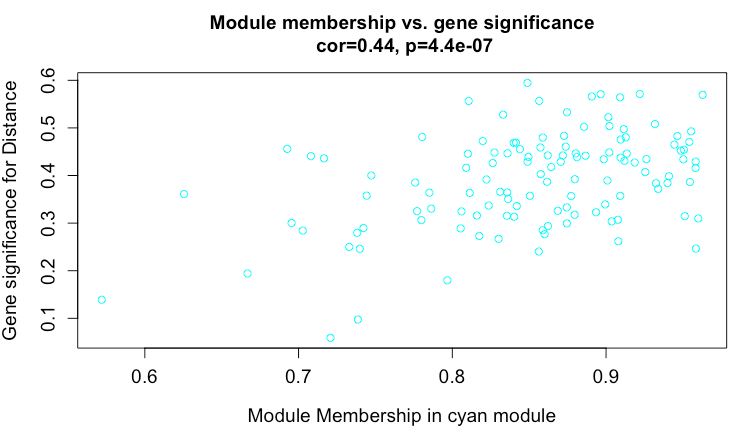

**Figure S7**. In the preoptic area of the hypothalamus, gene significance for distance to the decoy correlated with cyan module membership.

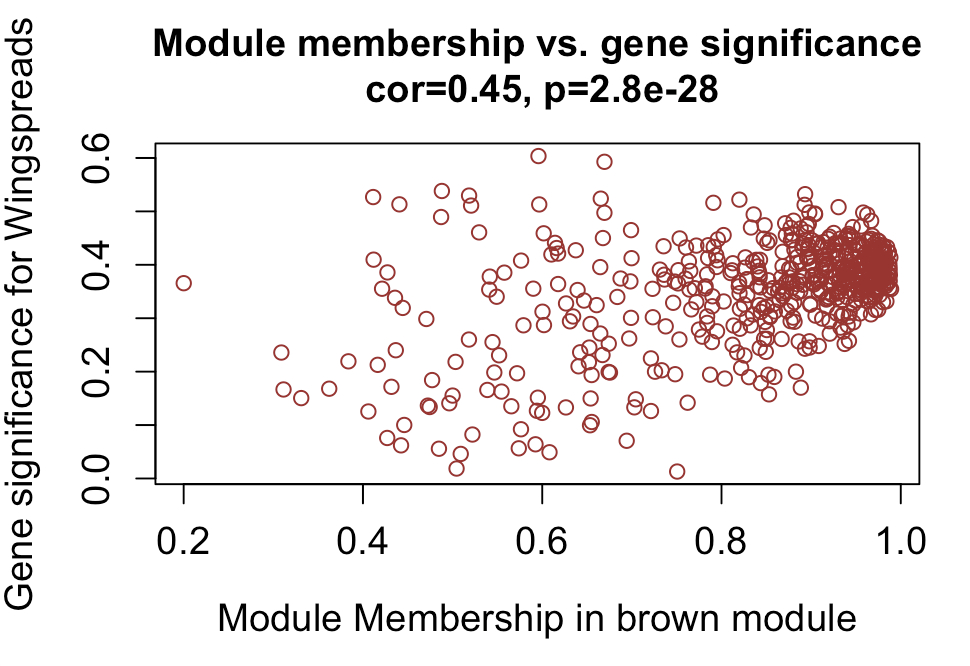

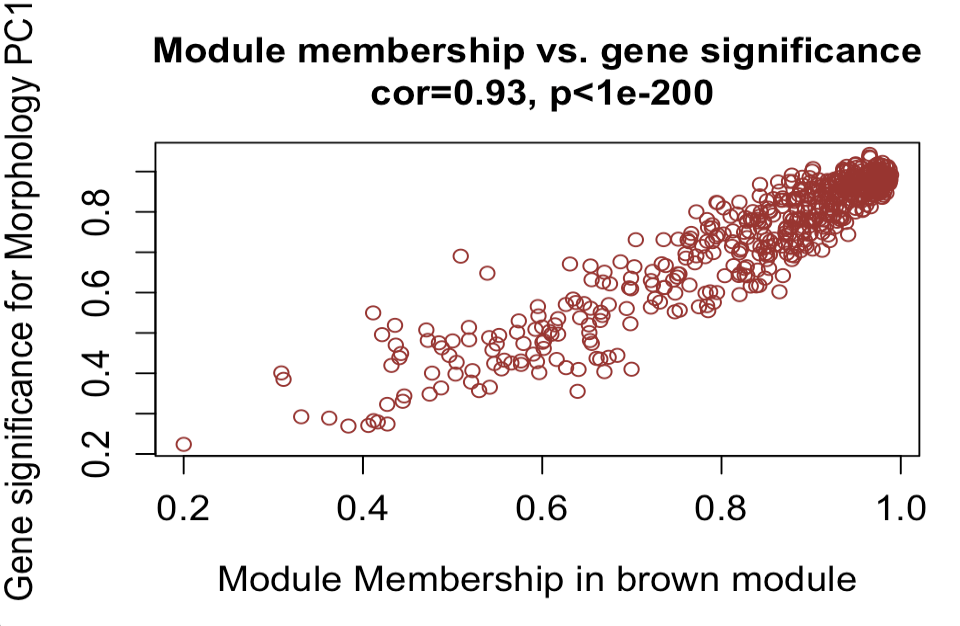

**Figure S8**. In the preoptic area of the hypothalamus, gene significance for A) wingspreads and B) morphology PC1 correlated with brown module membership.

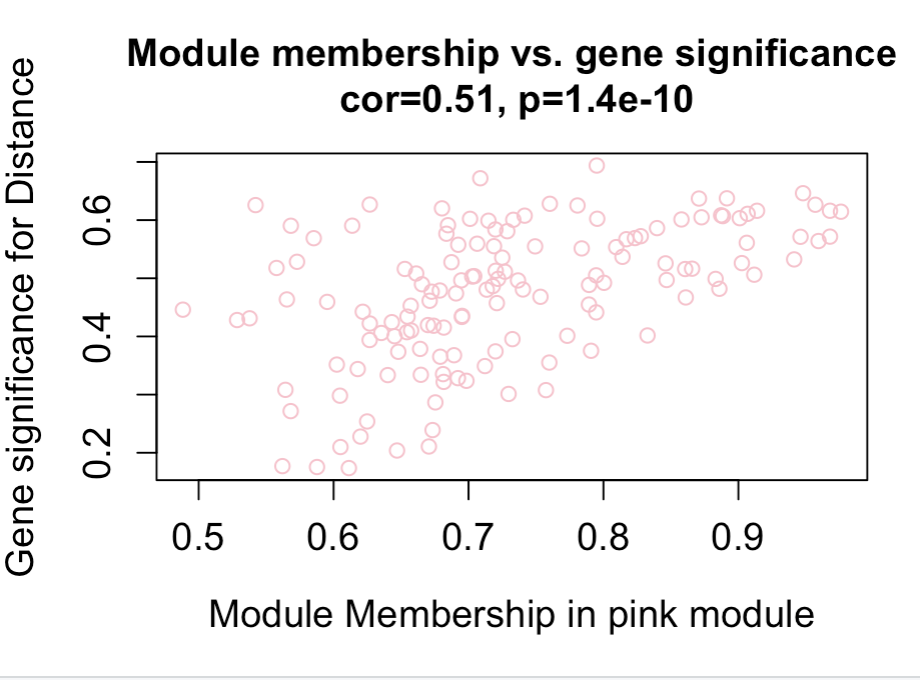

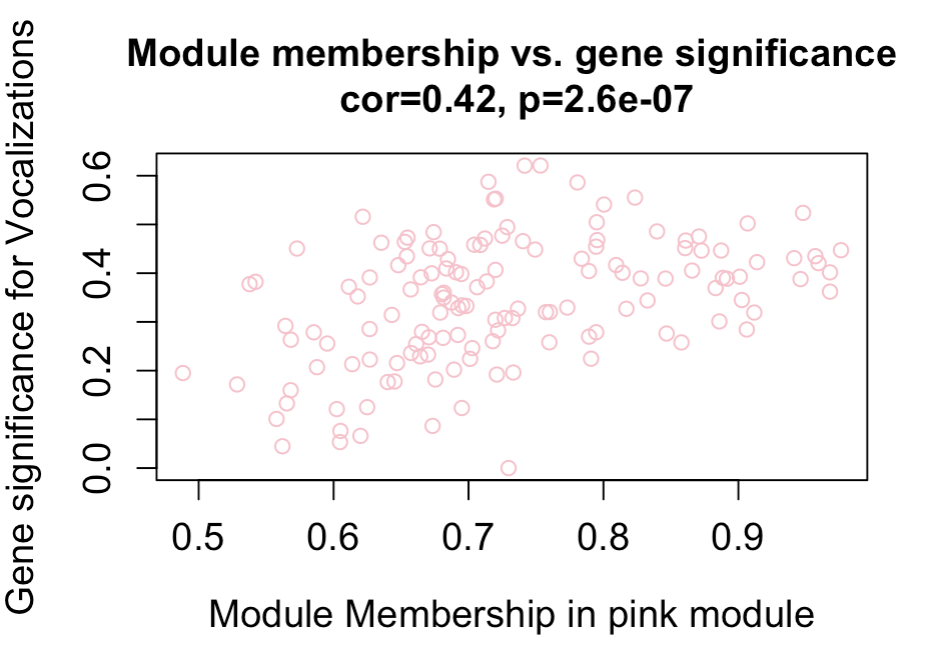

**Figure S9**. In the nucleus taeniae, gene significance for A) distance to the decoy and B) vocalizations correlated with pink module membership.

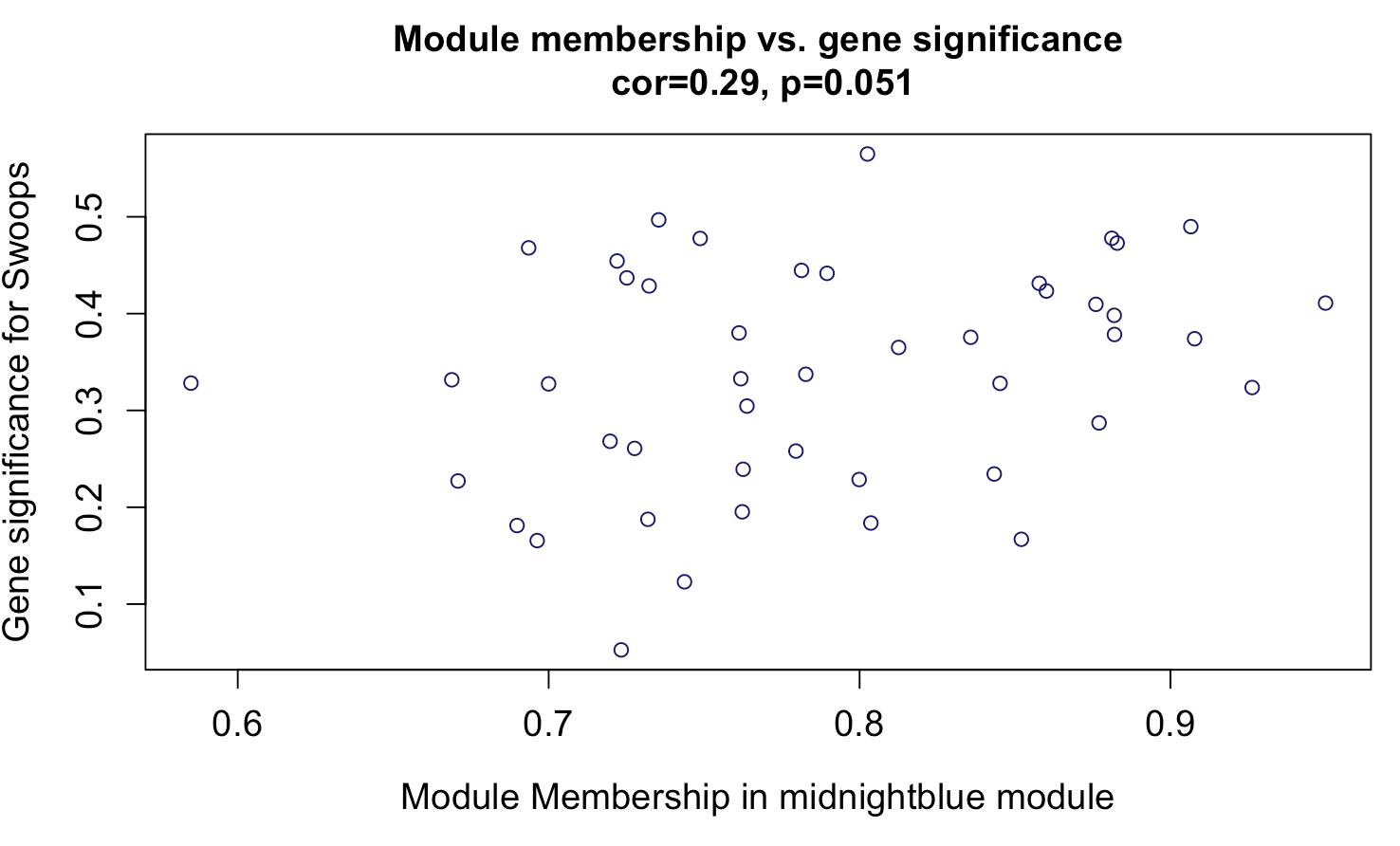

**Figure S10**. In the nucleus taeniae, gene significance for swoops correlated marginally with midnightblue module membership.

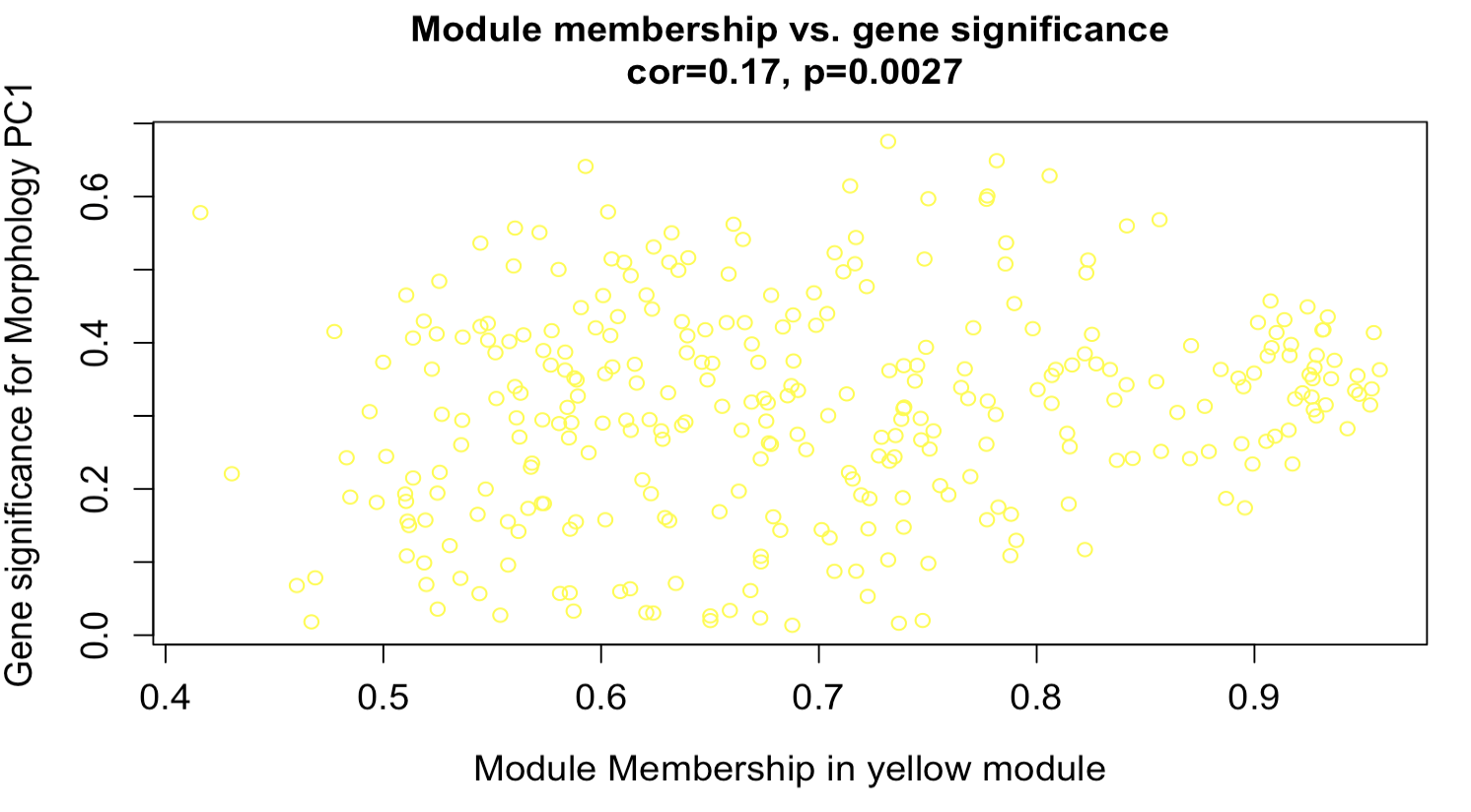

**Figure S11**. In the nucleus taeniae, gene significance for morphology PC1 correlated with yellow module membership.

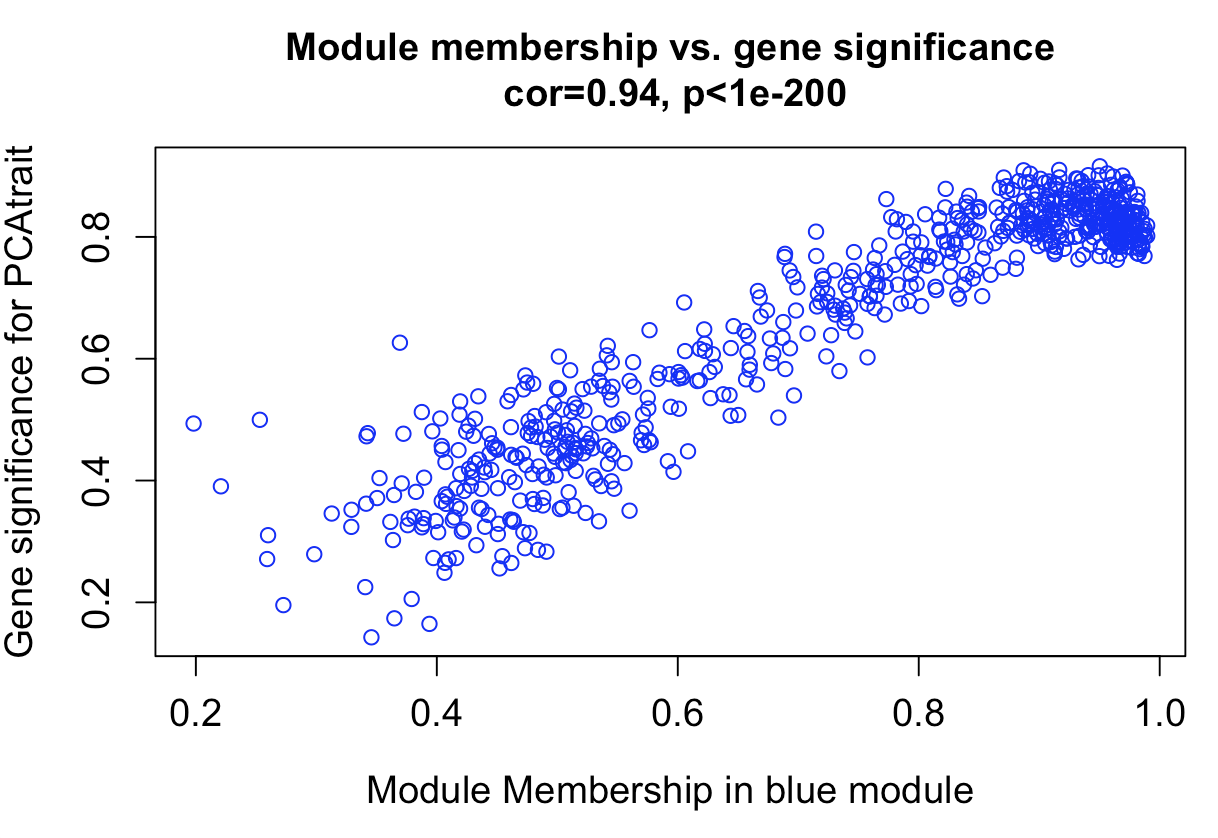

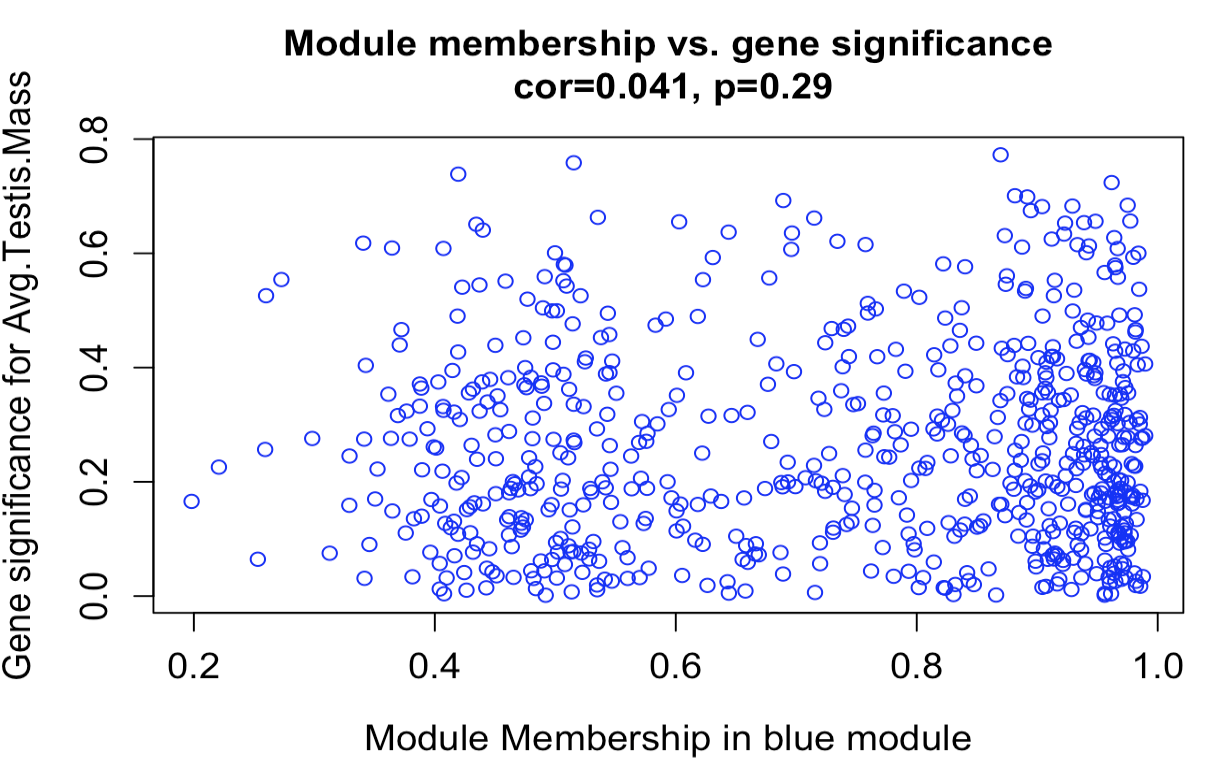

**Figure S12**. In the nucleus taeniae, gene significance for A) morphology PC1 correlated with blue module membership, but not for B) testes mass

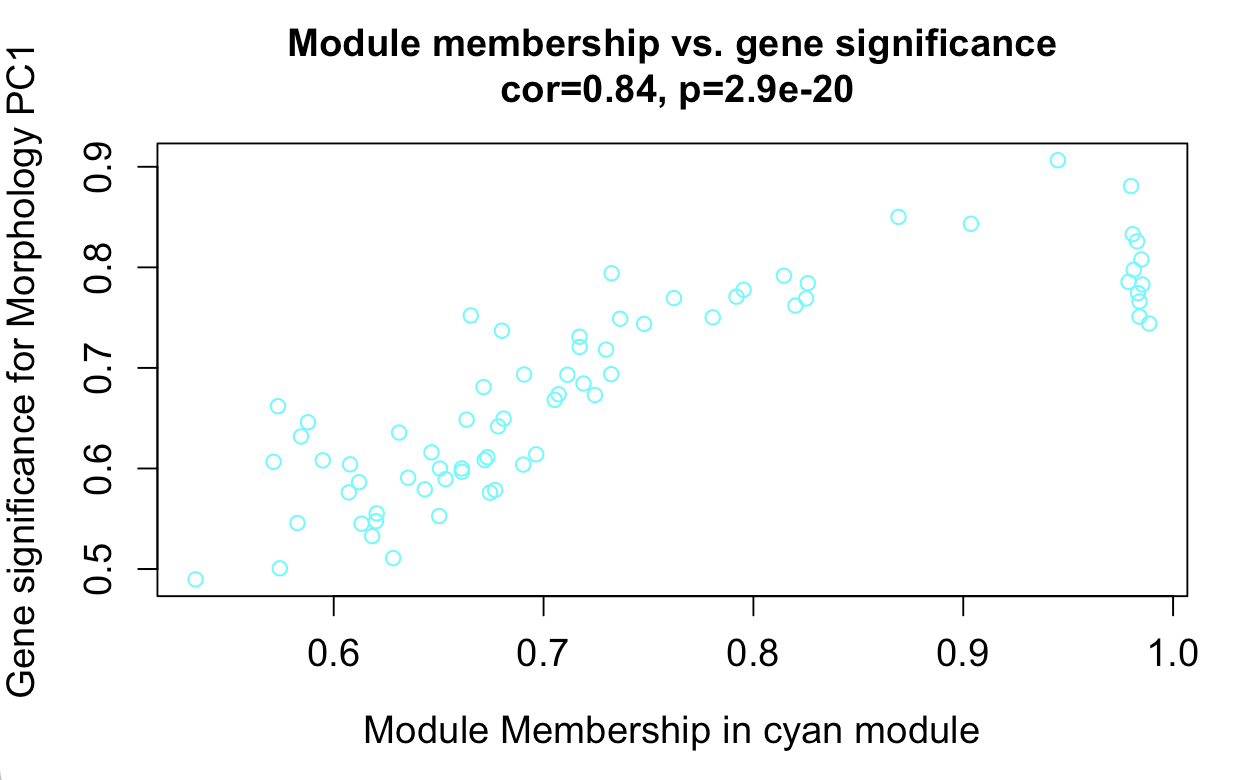

**Figure S13**. In the nucleus taeniae, gene significance for morphology PC1 blue module membership.

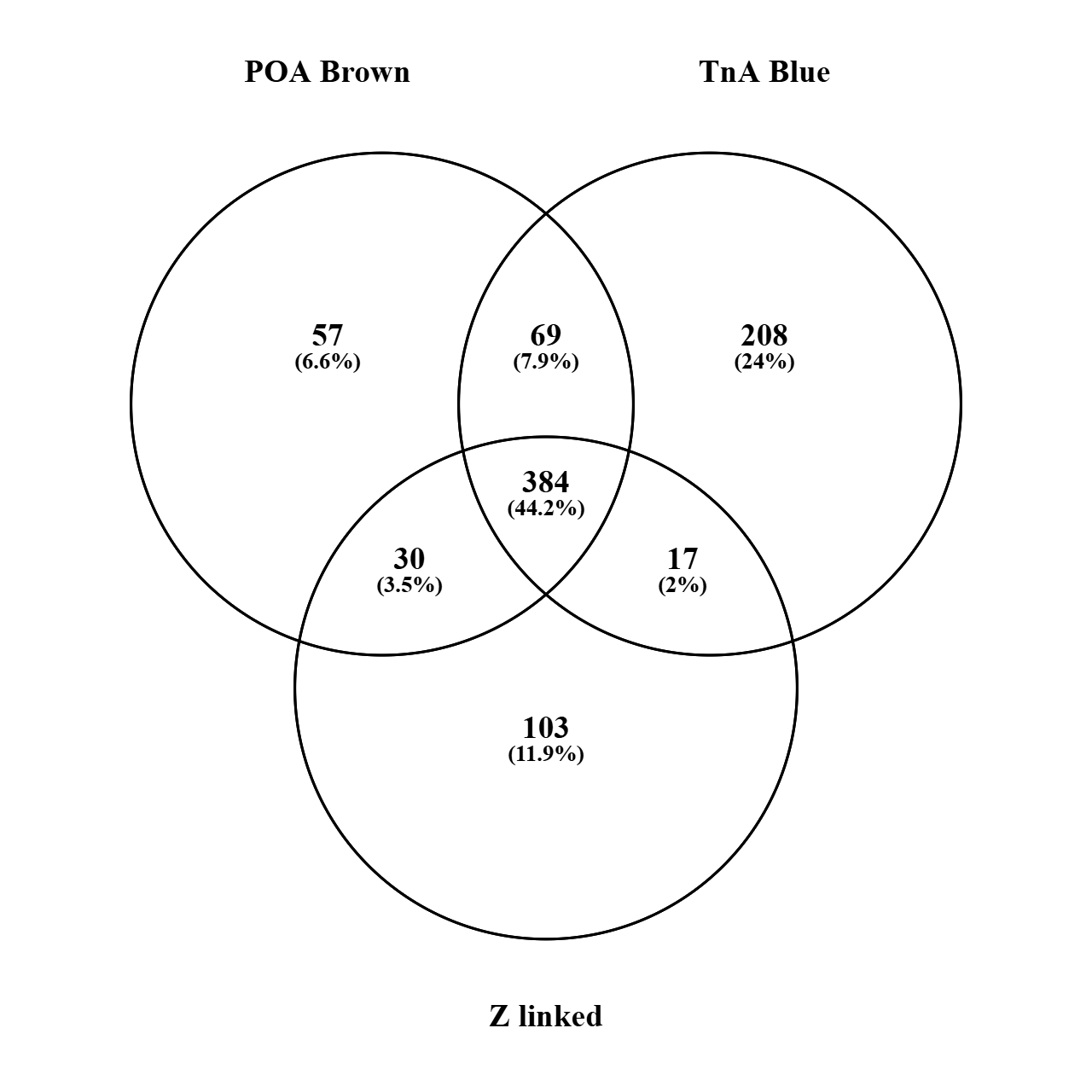

**Figure S14**. Venn diagram of genes shared between the blue module in the nucleus taeniae (TnA) and the brown module in the preoptic area of the hypothalamus (POA) with genes on the Z chromosome. The intersection between all three groups indicates that 384 Z-linked genes are shared between both modules.

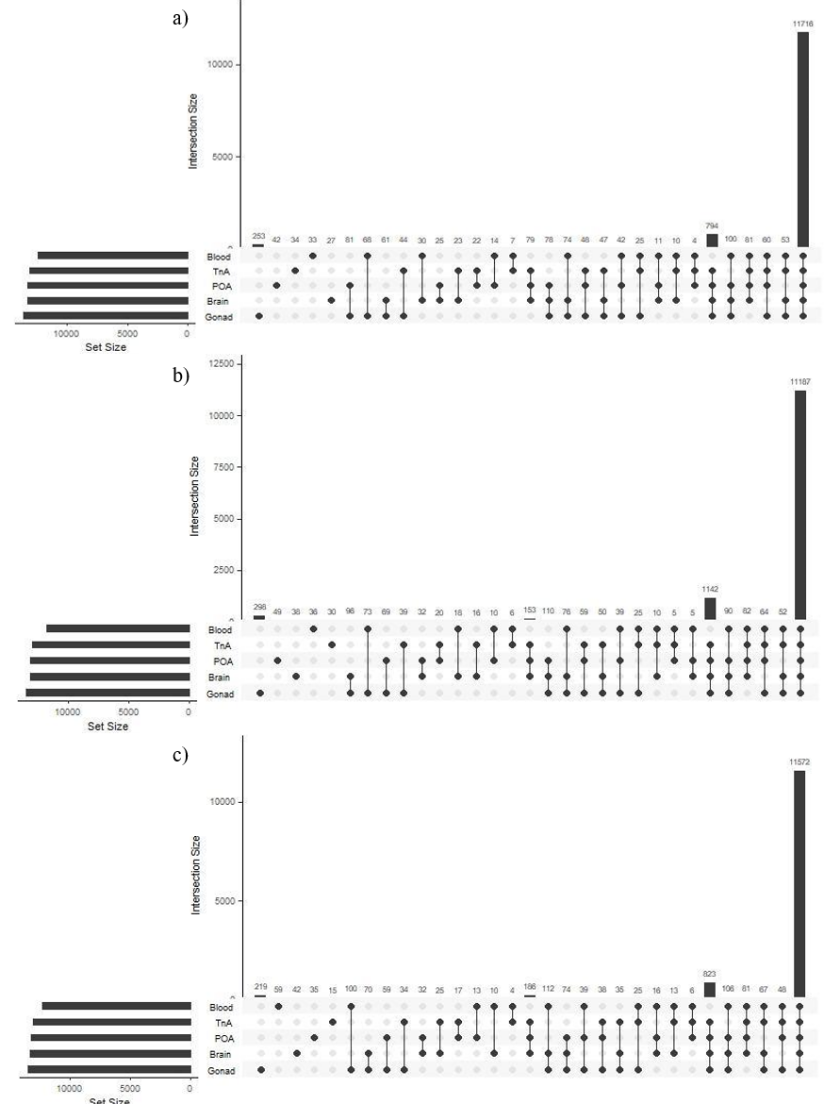

**Figure S15.** UpSet plot depicting the number of unique and shared genes in the whole blood, nucleus taeniae (TnA), preoptic area of the hypothalamus (POA), whole brain, and gonad from a) females, b) courting males, and c) parenting males. Intersection size is the number of shared genes present in these samples, and the black circles on the x-axis depict whether these genes are present or absent in that set.

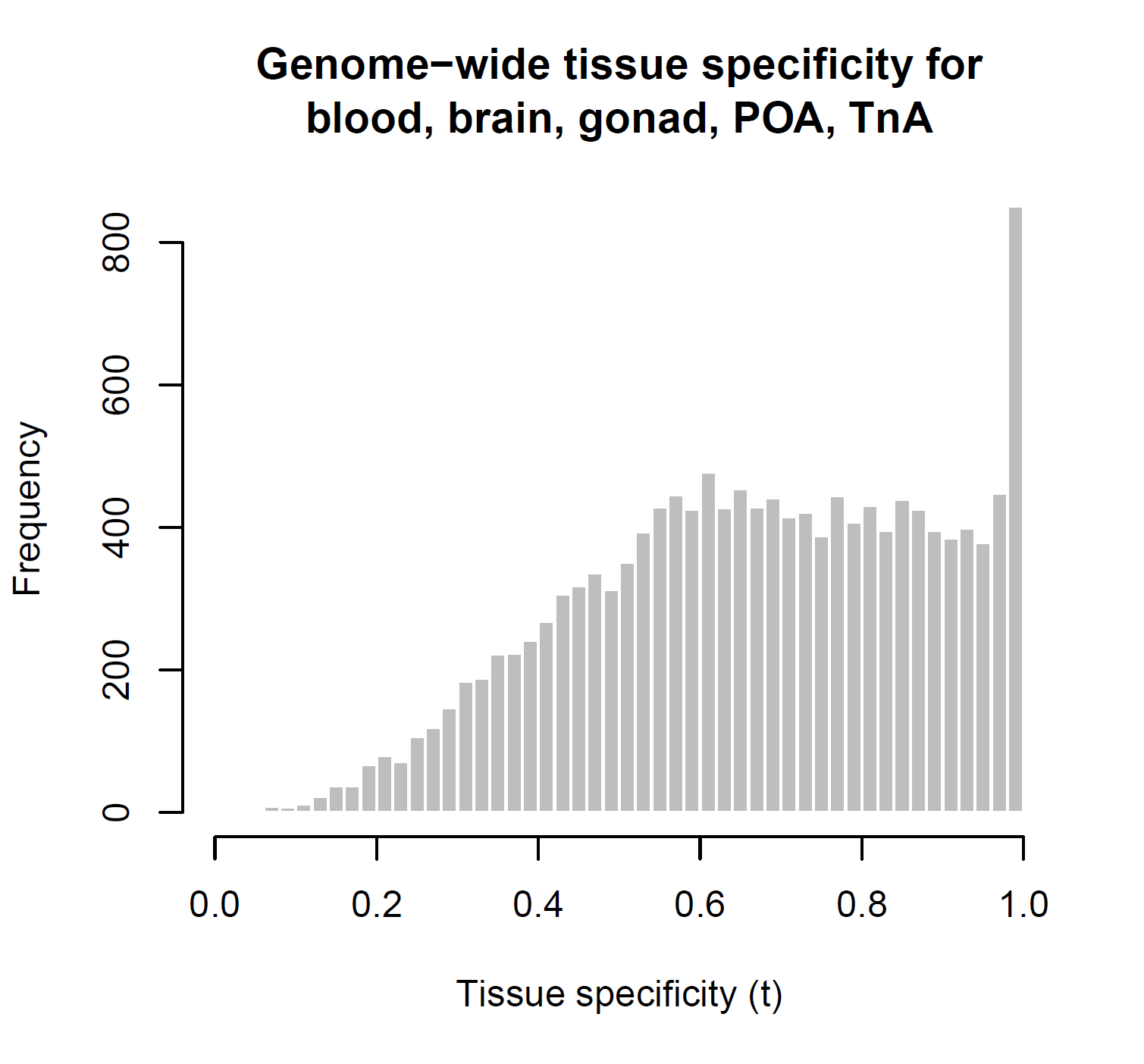

**Figure S16**. Tau index of gene tissue specificity (t) in blood, whole brain, gonad, preoptic area of the hypothalamus (POA), and nucleus taeniae (TnA).

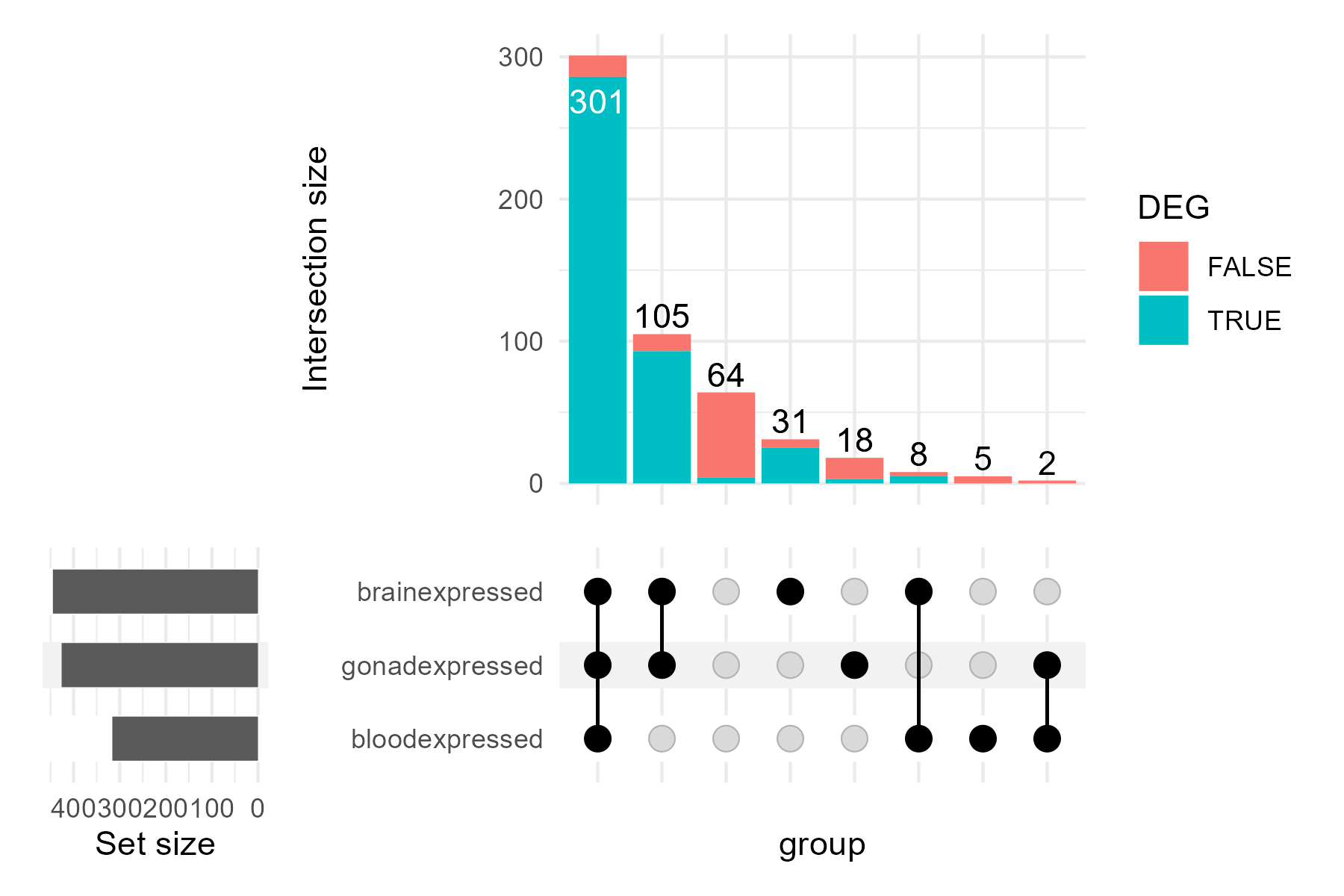

**Figure S17.** UpSet plot depicting the number of Z-linked genes expressed in whole brain, gonadal tissue, and blood. Intersection size is the number of shared genes present in these samples, and the black circles on the x-axis depict whether these genes are present or absent in that gene set. The blue genes in the intersection barplot represent genes that were differentially expressed in the preoptic area of the hypothalamus, nucleus taeniae, or both. Red genes were not differentially expressed. One intersection barplot (64) depicts genes that are absent from these tissues because these Z-linked genes are present in the jacana reference genome but were not expressed in these tissue samples. This intersection barplot also contains four differentially expressed genes that were found in either the preoptic area of the hypothalamus or the nucleus taeniae, but were not present in whole brain samples.
